# A yeast mating platform for multiplex screening of fungal GPCR-ligand interactions

**DOI:** 10.1101/2024.10.30.620951

**Authors:** Giovanni Schiesaro, Melani Mariscal, Mathias Jönsson, Ricardo Tenente, Mathies B. Sørensen, Marcus Wäneskog, María Victoria Aguilar-Pontes, Agustina Undabarrena, Marcus Deichmann, Emma E. Hoch-Schneider, Viji Kandasamy, Thomas M. Frimurer, Antonio Di Pietro, Line Clemmensen, Michael K. Jensen, Emil D. Jensen

## Abstract

Fungi are essential members across ecosystems, yet phytopathogenic fungi pose an increasing risk to crop yields. Despite their ecologic importance, cell-cell communication in fungi is underexplored, partly due to the lack of high-throughput techniques. Here we developed a Yeast Mating Platform (YeMaP) to investigate the interaction between fungal G protein-coupled receptors (GPCRs) and pheromone peptides. We used YeMaP for high-throughput screening of 8,000 pheromone sequences and identified new peptides with improved agonism or antagonism action. We found that these peptides can be applied in a native fungal system such as the plant pathogen *Fusarium oxysporum*, to control hyphal chemotropism and reduce plant root penetration. Additionally, we utilized YeMaP in a one-pot assay to investigate how abiotic factors influence the communication of multiple pheromone-GPCR combinations and found that the cell-cell communication mediated by the GPCR Ste2 from *F. oxysporum* signalled robustly across different abiotic factors, while other fungal GPCR-pheromone interactions were more sensitive to changes. Taken together, YeMaP accelerates the identification of fungal GPCR-peptide interactions by enabling one-pot assays, and serves as model system for studying fungal cell-cell communication.

## Introduction

Fungi are indispensable for fundamental processes of global ecosystems such as the cycling of carbon, nitrogen, and phosphorus, all of which are critical for the survival of plants and animals^1^. At the same time, pathogenic fungi pose an increasing threat to global food security^2^. As climate change intensifies, the available data and simulation models indicate that plant disease pressures are likely to rise substantially in the near future^3^. To address these emerging challenges, we need a greater understanding of how intra- and interspecies communication occurs in fungal consortia. G protein-coupled receptors (GPCRs) are master switches that convert external stimuli into an internally coordinated cellular response^4^. Through GPCRs, fungi can detect a variety of chemical signals including pheromones, nutrients, ions, hydrophobic surfaces, or light, allowing them to regulate their development, metabolism, and virulence^5^. However, our current understanding of the role of fungal GPCRs is limited^5^, and interactions between GPCRs and potential ligands are often poorly understood, thus hindering the development of new antifungal drugs and sustainable pest management systems for agriculture^6,7^.

The model yeast organism *Saccharomyces cerevisiae* encodes three GPCRs, two of which are mating receptors, *STE2* and *STE3* expressed in *MAT***a** and *MAT*α cells, respectively^8^. To form diploid cells, haploid *MAT*α cells communicate through an unmodified short peptide (α- factor) to *MAT***a** cells, which respond with a post-translationally modified peptide (**a**-factor)^9^. In contrast, in the asexual plant pathogen *Fusarium oxysporum (Fo),* autocrine pheromone communication regulates conidial germination^10^, and the *Fo STE2* homolog is required for hyphal chemotropism toward plant roots^11^. A similar mechanism has recently been shown in *Fusarium graminearum*^12^, pointing towards a conserved mechanism of plant recognition across the ascomycota phylum. Fungal mating receptors have been expressed in *Saccharomyces cerevisiae* for pathogen detection^13^ or to establish orthogonal communication^14^ in biosensor strains. However, cell-cell communication between GPCRs and pheromones has never been explored in multiplex assays to mimick the presence of multiple fungal organisms, although such a setup would better reflect the complexity of real- world interactions^15^. Additionally, fungal pheromones are partly conserved among closely related species^16^ and thus could function in interspecies communication^17^. This suggests that the yeast mating system could serve as a tool to study fungal cell-cell communication.

Unlike GPCR-based biosensors^18^, successful yeast mating requires the secretion and detection of pheromones along with the downstream activation of a developmental program for cell and nuclear fusion^19^. Here, we established a yeast mating platform (YeMaP) for analyzing interactions between fungal GPCRs and pheromone variants. We used YeMaP to study fungal cell-cell communication from four different perspectives: i) the interaction between a fungal GPCR and a pheromone, ii) screening of a library of peptide variants for identification of a synthetic pheromone with higher potency on the *F. oxysporum* Ste2 GPCR, iii) modeling cell-cell communication within fungal consortia, and iv) assessing the effects of external factors such as pH, nitrogen source, plant peroxidases, or different pheromone supplementations, on a network of library-on-library interactions. We further validate that both natural and artificial pheromones selected from YeMaP screening also confer function in the native fungal pathogen *F. oxysporum,* demonstrating that they trigger strong hyphal chemotropism while interfering with fungal penetration of plant roots. Altogether, YeMaP accelerates functional analysis of isolated fungal GPCRs, enabling large- scale exploration of GPCR-targeted pest management strategies.

## Results

### Engineering yeast mating to study fungal GPCR-pheromone interactions (One-on-One)

To define the best design for establishing a synthetic mating platform to study network interactions among multiple fungal GPCRs, we initially focused on the Ste2 GPCRs from *Candida albicans* (*Ca*) and *Fusarium graminearum* (*Fg*), which had previously been expressed in yeast^14,20,21^ (Fig. 1A). Since asymmetry in sexual pheromones is not required between yeast mating pairs^22^, we first sought to identify which combination of mating-types was more suitable for heterologous GPCR expression and alpha pheromone secretion^23^. To this aim, we co-cultured two yeast strains of opposite mating-types expressing either a fungal GPCR or a cognate pheromone, followed by growth-based diploid selection on plates^22,24^. We found that the substitution of the native *STE3,* with the heterologous GPCR, in *MAT*α cells and the production of the cognate alpha pheromone in *MAT***a** cells resulted in the highest diploid frequencies. Specifically, for the *Ca* GPCR-pheromone pair this setup yielded a 33-fold higher diploid frequency than the opposite combination (p<0.0001) (Fig. 1B), corroborating a previous study by Huberman *et al.*^24^.

**Fig. 1.**
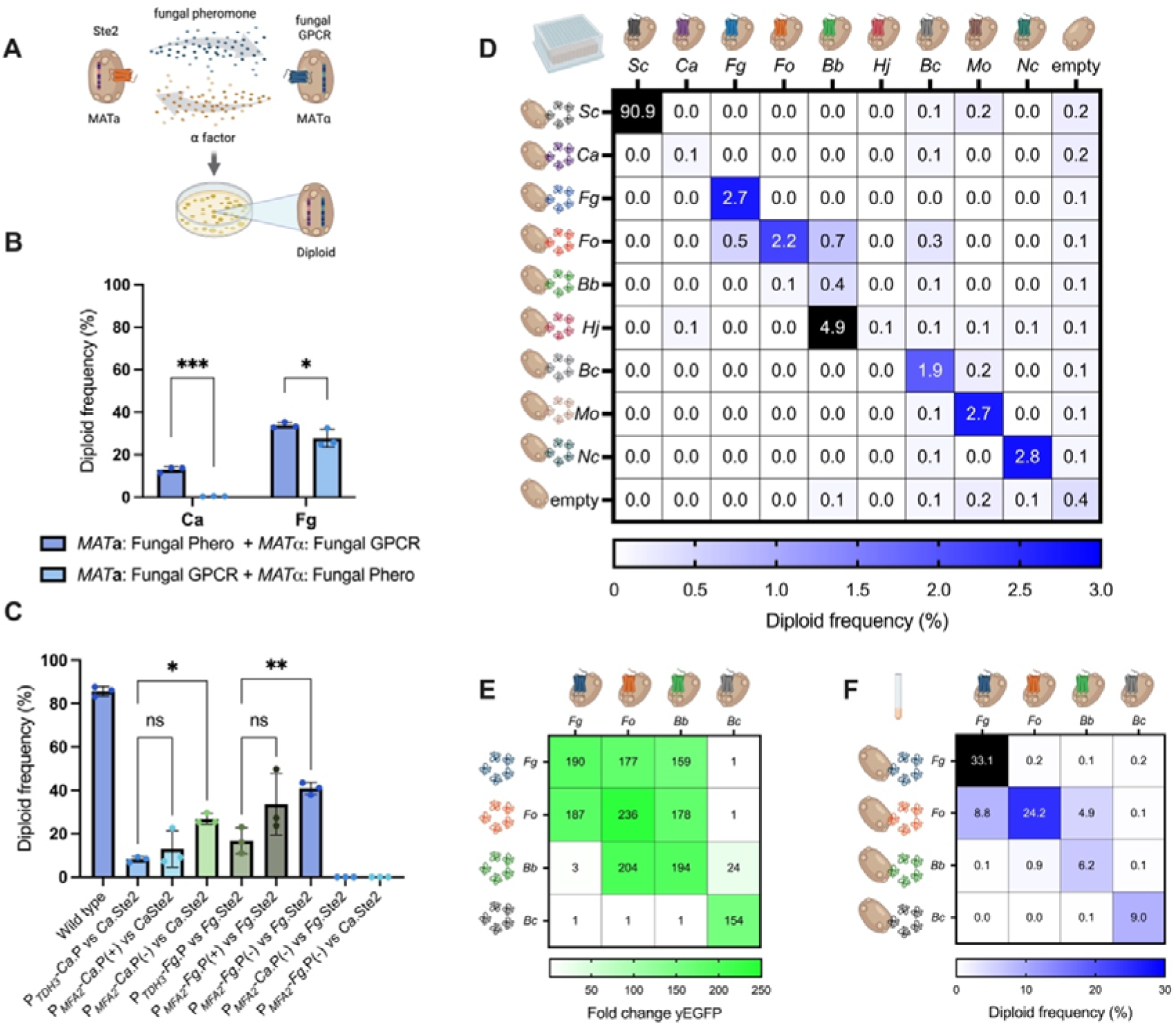
YeMaP correctly identifies fungal GPCR-pheromone interactions. **A** The One-on-One setting involves co-culturing two yeast strains in which half of the mating communication system has been replaced with heterologous fungal components; one cell expresses a given fungal Ste2 GPCR homolog while a second cell secretes a given fungal alpha pheromone. After successful mating, diploid cells are formed and selected. **B** Comparison of co-cultures expressing *Candida albicans* (*Ca*) or *Fusarium graminearum* (*Fg*) components under the following conditions: *MAT***a** cells expressing either the *Ca* or *Fg* alpha pheromone from the *MFA2* promoter (*Ca*: GEN29, *Fg*: GEN35) and *MAT*α cells expressing either the *Ca* or *Fg* Ste2 GPCR (*Ca*: GEN36, *Fg*: GEN37); or *MAT***a** cells expressing the fungal GPCR (*Ca*: GEN100, *Fg*: GEN78) and *MAT*α cells expressing the *Ca* or *Fg* alpha pheromone from the *MF*α*1* promoter (*Ca*: GEN76, *Fg*: GEN77). *MFA2* and *MF*α*1* promoters were selected as the strongest inducible pheromone promoters based on previous studies^20,29^. **C** Diploid frequency comparison between different designs. From left to right: GEN18 and GEN27 were co-cultured as a positive control (Wild type); GEN36 (*Ca*.Ste2) co-cultured with GEN28 (P*TDH3*-*Ca*.P); with GEN29 (P*MFA2*-*Ca*.P); or with GEN55 (P*MFA2*- *Ca*.P, *mfa1*Δ, *mfa2*Δ). GEN37 (*Fg*.Ste2) co-cultured with GEN34 (P*_TDH3_*-*Fg*.P); with GEN35 (P*_MFA2_*-*Fg*.P); or with GEN58 (P*_MFA2_*-*Fg*.P, *mfa1*Δ, *mfa2*Δ); and GPCRs combined with non-cognate pheromone GEN55 + GEN37; and GEN58 + GEN36. **D** Yeast mating matrix with *MAT*α strains individually expressing 8 different fungal Ste2 GPCRs (columns), co-cultured with either of 8 alpha pheromone-secreting *MAT***a** strains (rows). As positive and negative controls, respectively, strains expressing either *Sc* Ste2 or alpha pheromone, or strains lacking a GPCR or a pheromone were used. Blue gradient is set to a maximum of 3% of diploid frequency, with values >3% colored in black. Complete list of organisms and pheromone sequences is in Supplementary Table 1. **E** Activation levels of *Fg*.Ste2 (CPK366), *Fo*.Ste2 (GEN70), *Bb*.Ste2 (GEN71) and *Bc*.Ste2 (GEN128) biosensors (columns), incubated with 10 μM alpha pheromone from *Fg*, *Fo*, *Bb*, or *Bc* (rows). The fluorescent signal was normalized to the yEGFP background value (receptor without pheromone). **F** Validation of the mating matrix in culture tubes. Strains individually expressing the 4 indicated GPCRs (columns) were co-cultured with those secreting the indicated alpha pheromone (rows). Blue gradient is set to a maximum of 30% of diploid frequency, with values >30% colored in black. Plating was performed in three technical triplicates in **D**. Co-cultures were performed in three biological replicates and three technical replicates in **B**, **C,** and **F**. Experiments were conducted in four biological replicates in **E**. Statistical significance was determined in **B** through two-way analysis of variance (ANOVA) with Tukey’s multiple comparisons (✱p□≤□0.05, ✱✱✱p□≤□0.001); and in **C** with one-way ANOVA with Tukey’s multiple comparisons tests in GraphPad Prism (✱p□≤□0.05, ✱✱p□≤□0.01).

Next we asked if mimicking increased pheromone expression during native yeast mating is an essential feature^9,20^. This was done by comparing diploid frequencies from strains showing either constitutive or inducible expression of the cognate heterologous pheromone. Higher diploid frequencies were obtained when the heterologous pheromones were expressed from the inducible *MFA2* promoter^25^ compared to the strong constitutive *TDH3* promoter^26^, although the difference was only statistically significant when the strains were deleted for both **a**-factor genes, *MFA1* and *MFA2* (p=0.039 with *Ca*.Ste2 and p=0.005 with *Fg*.Ste2) (Fig. 1C). Importantly, no diploid formation was observed when a given GPCR- expressing strain was co-cultured with strain expressing a non-cognate pheromone (Fig. 1C).

Once the optimal strain design had been established, we evaluated the specificity of the YeMaP by performing a “One-on-One” mating matrix with strains expressing 8 different fungal GPCR and strains secreting 8 different pheromones. The native *Sc* system and an empty construct were added as positive and negative controls, respectively (Fig. 1D). Here we observed promiscuity of the *Beauveria bassiana* (*Bb*) GPCR (*Bb*.Ste2) towards pheromones from other species such as *F. oxysporum (Fo)* or *Hypocrea jecorina* (*Hj).* Furthermore, *Fo* pheromone induced diploid formation upon co-culture with cells expressing the GPCRs *Fg*.Ste2, *Fo*.Ste2, or *Bb*.Ste2 (Fig. 1D).

We performed further validation in the biosensor assay^14^ with four selected GPCRs (*Fg*.Ste2, *Fo*.Ste2, *Bb*.Ste2, and *Botrytis cinerea* (*Bc*) *Bc*.Ste2) and their chemically synthesized cognate pheromones. Biosensor assays^14^ were performed using a Green Fluorescent Protein (yEGFP) reporter as readout (Fig. 1E), while in parallel comparing the results to the mating-based selection assay in culture tubes, to evaluate which of the two approaches yielded the highest resolution (Fig. 1F). We observed a binary (on/off) behavior for the biosensors, with *Bb*.Ste2 showing almost full activation in response to *Fg* pheromone, and *Fo*.Ste2 exhibiting a strong activation when exposed to either *Bb* or *Fg* pheromones (Fig. 1E). However, these three strong heterologous GPCR-ligand responses did not correlate with diploid formation rate in the Yeast Mating Platform (YeMaP), which were lower in comparison to the corresponding cognate pheromone interactions (Fig. 1F). The observed differences between mating pathway activation in the biosensor assay and diploid formation in YeMaP suggests that pheromone-induced signaling alone is not sufficient to trigger the much more complex mating response. Validation of the mating matrix in culture tubes recapitulated the same trends as shown in Fig. 1D, yet with a higher diploid frequency in this condition (Fig. 1F).

To further explore on the observed cross-reactivity between *Bb*.Ste2 and *Fo* pheromone detected in the YeMaP assay, along with the documented protective effect of *Bb* against *Fo* on crops^27,28^, we investigated if the species *Bb* is able to sense and respond to *Fo* pheromone. Indeed, either *Bb* or *Fo* pheromone induced a significant chemotropic response in *Bb* germ tubes (p<0.05) at a concentration of 378 μM, whereas a scrambled version of the *Fo* pheromone did not (Supplementary Fig. 1).

Taken together, these results demonstrated that the YeMaP assay provides a superior resolution in deciphering GPCR-pheromone interactions than the previously reported yeast biosensor assay. We therefore asked if YeMaP, unlike biosensors, is able to scale investigations by enabling mating-based selection of pheromone-GPCR interactions in a multiplex setup.

### Screening a library of pheromone variants on a fungal GPCR to identify improved pheromone agonists (Library-on-One)

We next asked whether the YeMaP assay could be used to screen for improved alpha pheromone agonists. Several *Sordariomycetes* with relevant ecological roles have alpha pheromones sharing a similar backbone^16,30^. Given this natural resemblance, and building on the deciphered interactions shown above (Fig. 1F), we constructed an alpha pheromone library (named YPL1) encoding 8,000 pheromone peptide variants to reach saturation of three internal residues in the sequence WCXXXGQPCW (where X is any possible amino acid in position 3, 4, and 5), while preserving the two essential residues Gly6-Gln7 responsible for maintaining the three-dimensional structure of the *Fo* alpha pheromone^31^. By adapting a recently established state-of-the-art yeast transformation method^32^, we transformed over 1.8 x 10^6^ yeast cells, obtaining a 6.9-fold coverage of our DNA-encoded pheromone library (8,000 peptides = 262,144 nucleotide variants), with each individual cell expressing a single alpha pheromone variant.

To search for pheromone agonists in a multiplex pool, we co-cultured the YPL1 library with cells individually expressing Ste2 GPCRs from the plant pathogens *Fg*, *Fo*, or *Bc*, or from the insect pathogen *Bb*. The culture was sequentially enriched for diploid cells before the DNA was extracted from the pool and analyzed by Next-Generation Sequencing (NGS) (Fig. 2A). After normalizing the counts in Reads Per Million (RPM) and subtracting the reads of the library before the co-cultures, we observed a specific and exclusive enrichment for the *Bb* alpha pheromone in the *Bb*.Ste2 co-cultures (+307 RPM) and for the *Fg* alpha pheromone in the *Fg*.Ste2 co-cultures (+807 RPM). Strikingly, the *Fo* alpha pheromone was by far the most highly enriched sequence for the four Ste2 receptors tested in our screen (>340,000 RPM per sample).

**Fig. 2.**
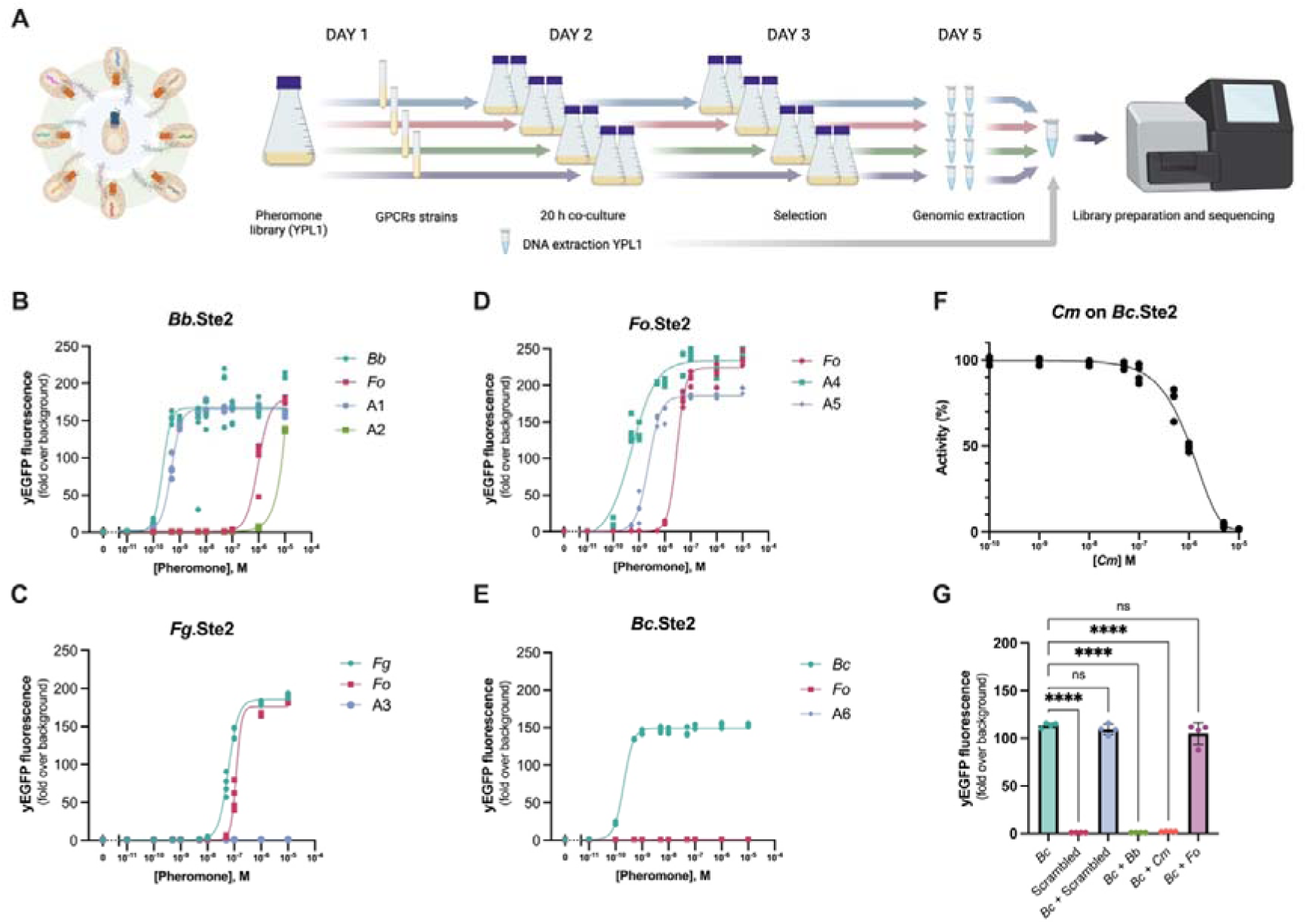
Identification of agonists and an antagonist by screening a library of pheromone variants. **A** A library of 8,000 alpha pheromone variants expressed in yeast (YPL1) was co-cultured with yeast strains expressing GPCRs *Bb*.Ste2 (GEN88), *Fg*.Ste2 (GEN87), *Fo*.Ste2 (GEN90) or *Bc*.Ste2 (GEN89) for 20 h in shake flasks (2 flasks per GPCR). After 2 days of enrichment for diploid cells, gDNA was extracted and used for sequencing on NextSeq 500. **B-E** Dose-response curves of the most enriched pheromones in combination with **B** *Bb*.Ste2 (*Fo*, A1 and A2); **C** *Fg*.Ste2 (*Fo* and A3); **D** *Fo*.Ste2 (*Fo*, A4, and A5); and **E** *Bc*.Ste2 (*Fo* and A6) were compared with the cognate pheromone of each GPCR (*Bb*, *Fg*, *Fo*, and *Bc*, respectively). **F** Inhibition curve of *Bc*.Ste2 (GEN128) co-cultured with 0.0005 μM of cognate *Bc* pheromone and increasing concentrations of *Cm* pheromone. **G** Fold change activation of *Bc*.Ste2 over non-treated control upon addition of 0.0005 μM *Bc* pheromone alone or in combination with 10 μM of either Scrambled (negative control), *Bb*, *Cm* or *Fo* pheromone. 10 μM of Scrambled peptide alone was used as negative control. Note that *Bc* + Scrambled and *Bc* + *Fo* do not differ significantly from *Bc* alone, and that activation by *Bc* was completely abolished in the presence of *Bb* or *Cm* pheromones. Means represent four biological replicates. Statistical significance was determined using one-way ANOVA with Dunnett’s multiple comparison test in GraphPad Prism (✱✱✱✱*p*□<□0.0001).

To validate the YeMaP results, we performed dose-response curves with chemically synthesized versions of 10 different alpha pheromone sequences selected among the top 20 most abundant pheromone sequences obtained from both biological replicates for each GPCR (Supplementary Fig. 2 and Supplementary Table 2). When incubated with the Ste2 GPCR for which they were originally enriched, seven out of the ten selected peptides induced strong activation (≥150-fold increase over background) of the pheromone response pathway (Fig. 2B-E), while two out of the remaining three pheromones induced a low activation (≥1.5-fold increase over background) (Supplementary Fig. 3A).

The *Bb* and *Fo* alpha pheromones, which previously produced similar diploid frequencies (Fig. 1D), displayed two distinct half-maximal effective concentration (EC_50_) values in the *Bb*.Ste2 biosensor strain (Fig. 2E). The EC_50_ of the *Fo* pheromone was 3,910-fold higher than that of the Bb pheromone (*Fo*_EC_50_ = 9.58 x 10^-7^ M and Bb_EC_50_ = 2.45 x 10^-10^ M). We further noted that the alpha pheromone Agonist 1 (A1), which contains a conserved Arg4 residue preceded by Leu3 and followed by Pro5, a combination not found in any known organism (Supplementary Fig. 2), exhibited a potency similar to that of the *Bb* pheromone (A1_EC_50_ = 4.82 x 10^-10^ M). Furthermore, our screening identified two other agonist pheromones, A4 and A5, both carrying a conserved Trp4, whose EC_50_ for *Fo*.Ste2 was at least 13-fold lower than that of the native *Fo* alpha pheromone (A4_EC_50_ = 5.16 x 10^-10^ M, A5_EC_50_ = 2.31 x 10^−9^ M, compared to Fo_EC_50_ = 2.92 x 10^-8^ M). While the A5 sequence was present in the *Fo* alpha factor pro-sequence (Supplementary Table 4), the A4 agonist contains an Ala5, representing a novel synthetic alpha pheromone with significantly increased potency, which requires 57 times less pheromone to achieve the same EC_50_ as the native *Fo* alpha pheromone (Fig. 2D).

Molecular dynamic (MD) simulations suggested that *Fo* pheromone, A5, and A4 share the same binding pose within the orthosteric pocket of *Fo*.Ste2, in which the carboxylic acid functional group of Trp10 forms salt bridges with Arg84 and Arg187 (Supplementary Fig. 4). We further noted that A4 had the most stable conformation from residues 5-10, especially in Trp10, which is the anchoring residue to the bottom of the orthosteric binding pocket (Supplementary Fig. 4B).

Next, we asked if our YeMaP dataset could also detect potential alpha pheromone antagonists. By searching for underrepresented sequences in our analysis, we noted that a peptide encoded in the alpha factor pro-sequences of the entomopathogenic fungi *Cordyceps militaris* and *Beauveria asiatica* (Supplementary Table 4) was consistently lacking in the *Fg*.Ste2, *Fo*.Ste2, and *Bc*.Ste2 post mating enrichments, while its enrichment was observed for *Bb*.Ste2 (+7,508 RPM). Therefore, we speculated that this alpha pheromone (named *Cm*) could function as a potential GPCR antagonist that inhibits diploid formation. To test this idea, we obtained the chemically synthesized *Cm* pheromone and found that it triggered the activation of *Bb*.Ste2 and *Fo*.Ste2, but not of *Fg*.Ste2 or *Bc*.Ste2. When co-incubated with the lowest pheromone concentration capable of inducing GPCR activation, the *Cm* pheromone showed an antagonistic effect on *Bc*.Ste2 (Supplementary Fig. 3B), with a half-maximal inhibitory concentration (IC_50_) of 1 μM (Fig. 2F). This IC_50_ was reached with a 2,000-fold excess of the antagonist compared to the agonist. A similar antagonistic effect was detected for the *Bb* pheromone during co-incubation with *Bc*.Ste2, while no such effect was observed with the *Fo* pheromone (Fig. 2G). Interestingly, the two antagonistic pheromones *Cm* and *Bb* share eight out of nine residues with the *Bc* pheromone. Furthermore, the antagonistic effect was only observed when the two residues Arg4 and Pro5 (not present in *Fo* pheromone) were preceded by either Leu3 or Met3 instead of Gly3, combined with the presence of an additional Trp10 residue, which is lacking in the nine-residues *Bc* pheromone (Supplementary Tables 1-2). The antagonistic effect of the *Cm* pheromone was further confirmed in a mating assay. When a co-culture consisting of strains expressing *Bc* GPCR and *Bc* pheromone was incubated with 50 μM of *Cm,* a 2-fold reduction in diploid formation (p=0.0071) was detected compared to the same co-culture incubated with 50 μM of scrambled peptide (Supplementary Fig. 3D).

In conclusion, by using a straightforward "Library-on-One" YeMaP assay we identified novel alpha pheromone agonists as well as an antagonist, by choosing the most or least enriched sequences, respectively, followed by simple validation assays.

### Abiotic and external factors shape the interaction networks between multiple fungal pheromone-GPCR signalling components (Library-on-Library)

External parameters such as pH or nitrogen source act as potent regulators of fungal growth, development, and pathogenicity^33^. Here we analyzed the effect of such abiotic factors on fungal GPCR-based cell-cell communication in multiplex (Fig. 3A). To this aim, we introduced a barcode system based on the Cre-lox recombinase^34^, which allows to genetically detect the occurrence of mating between two yeast strains, each expressing a given Ste2 GPCR or an alpha pheromone. In brief, each haploid cell contains a conserved region flanked by a unique barcode of 20 bp, a spacer, and a Lox site (Table 5). After successful mating, a unidirectional recombination event positions the two barcodes on the same chromosome^35^ (Fig. 3A). Quantitative PCR (qPCR) was used to detect the two barcodes and reveal the association between any given pheromone and GPCR, allowing us to estimate the relative abundances of diploid cell type in a pool. The relative abundance of a diploid combination was compared with the abundance of all diploids containing the conserved regions (R1 + R3) flanking the two barcodes (see Methods section).

**Fig. 3.**
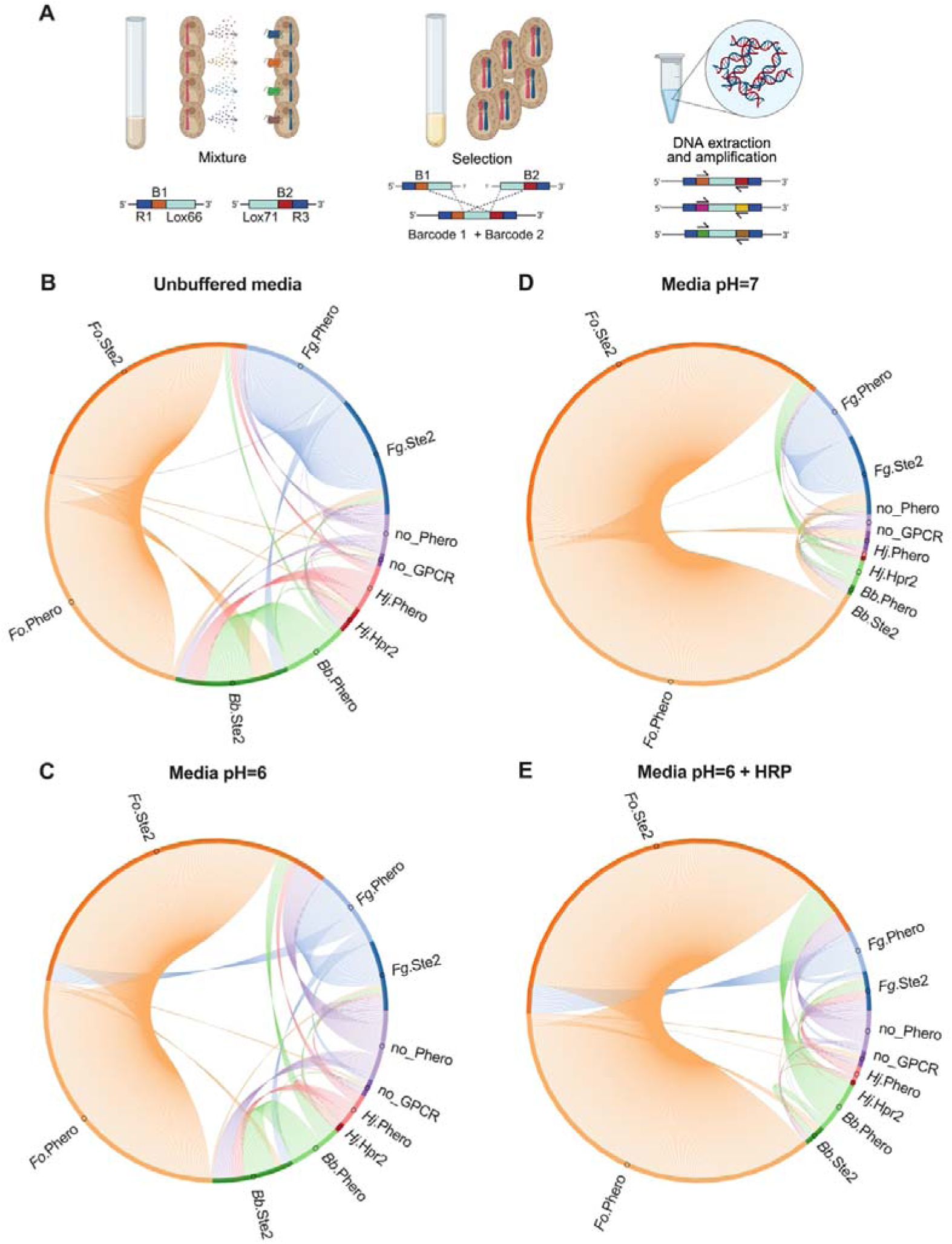
Abiotic and external factors govern fungal cell-cell communication mediated by GPCR-pheromone interactions. **A** Schematic illustration of the barcode system. *MAT***a** and *MAT*α cells contain a barcode flanked by a conserved region and a Cre-Lox recombination site on chromosome X. After successful mating the two barcodes can recombine and become located on the same chromosome. **B-E** Chord diagrams of consortia combining yeast strains expressing either *F. graminearum* (*Fg*), *F. oxysporum* (*Fo)*, *H. jecorina* (*Hj*), *B. bassiana* (*Bb*) Ste2 GPCRs with strains individually secreting the respective cognate alpha pheromones, as well as the two negative controls (empty strains) in the following media: **B** Unbuffered pH=5.6; **C** buffered at pH=6; **D** buffered at pH=7; and **E** buffered at pH=6 supplemented with 1 μM Horse Radish peroxidase (HRP). Dark color represents the GPCR-expressing strain while the lighter version represents the strain secreting the cognate alpha pheromone. Orange=*Fo*, Blue=*Fg*, Green=*Bb*, Red=*Hj*, Violet=negative control with empty constructs. Arcs represent the connection between a GPCR and a pheromone-expressing strain. The size of an arc reflects the average relative abundance of a diploid cell. Relative quantification was done by comparing the cycle threshold (Ct) value of every possible combination of two barcodes with the Ct value of the conserved region flanking all barcodes. All experiments were conducted in three biological replicates.

We first tested the specificity of the system by running 4 different co-cultures and combinations of these co-cultures in consortia, by assessing whether the two barcode sites were on the same chromosome (Supplementary Fig. 5). The strains lacking constructs (no_Pheromone, no_GPCR) were capable of forming diploids in co-culture (Fig. 1D and Supplementary Fig. 5), but were not detected in consortia with other strains (Supplementary Fig. 5). Introduction of barcodes in the remaining strains expressing pheromones or GPCRs allowed specific amplification and quantification of diploids by qPCR. From all co-cultures tested, the relative abundance of diploids detected with each set of primers was comparable (Supplementary Fig. 6 and Supplementary Data 3).

To mimic the presence of multiple fungal organisms of ecological interest, we performed consortia combining yeast strains expressing either *Fg*, *Fo*, *Hj*, *Bb* Ste2 GPCRs with strains individually secreting the respective cognate alpha pheromones, as well as the two negative controls (empty strains). These 10 different strains were combined into pre-culture tubes under different conditions, including three different pHs (unbuffered, pH=6, or pH=7), or in the presence of Horse Radish Peroxidase (pH=6 + HRP) to mimic a plant root environment^11^. Interestingly, the *Fo* pheromone-GPCR system was the most robust and consequently more effective in the formation of diploid cells in all the conditions tested (Fig. 3B-E). Conversely, in the *Fg* pheromone-GPCR system which had the second most abundant diploid formation in unbuffered media or in media buffered at pH=6 or pH=7, the presence of HRP significantly reduced the formation of diploid cells (p=0.023, Supplementary Fig. 7A).

Overall, with the exception of *Fo* GPCR and *Fo* pheromone-expressing strains, a reduction in diploid formation was observed at increasing pH values or in the presence of HRP (Supplementary Fig. 7A). Importantly, the results of this library-on-library approach were consistent with those from individual co-cultures (one-on-one) containing only one strain expressing a given GPCR and another secreting a given alpha pheromone from *Fg*, *Fo*, *Bb*, or *Hj* (Supplementary Fig. 7B).

In contrast to the above approach, the yeast biosensor assay failed to detect a difference in signaling behaviour between *Fo*.Ste2 and the remaining GPCRs when exposed to HRP. We observed activation of the control strain lacking a GPCR in unbuffered Synthetic Complete (SC) media supplemented with 2 μM HRP, as well as a general activation (≈1.5 fold increase in yEGFP fluorescence) at pH=6 with 2 μM HRP, both in the control strain and in all the biosensors tested (Supplementary Fig. 8A-B).

Taken together, these results demonstrate that a synthetic mating platform can be used to simultaneously study the effect of abiotic factors on cell-cell communication between fungal GPCRs and alpha pheromones. Furthermore, we found that a library-on-library approach leads to similar results as a one-vs-one approach, and that *Fo* GPCR-pheromone communication was not affected by the different external factors tested.

### External pheromone supplementation alters the cell-cell communication of phytopathogen GPCRs in a fungal consortium

After using the YeMaP platform to determine the effect of abiotic factors on four different fungal GPCR-pheromone pairs and showing the robustness of the *Fo* signalling components, we next explored if external addition of a pheromone could reshape a network of pheromone-GPCR interactions. With the goal of limiting the pheromone-GPCR systems from the two plant pathogens *Fo* and *Fg*^36^ and promoting those from the non-pathogenic/commensal fungi *Bb* and *Hj^37,38^*we tested the effect of supplementing either scrambled, *Cm*, or A4 peptide.

In SC media without external pheromone supplementation or containing 10 μM of scrambled peptide, almost all diploids detected were formed from the *Fo*.Ste2 + *Fo*.Pheromone or *Fg*.Ste2 + *Fg*.Pheromone pairs (Fig. 4A-B). By contrast, external supplementation of *Cm* pheromone resulted in a 0.2-fold decline in diploids containing *Fo*.Ste2 (p=0.001) concomitant with a 2.2-fold increase in diploids containing *Fg*.Ste2 (p=0.015), while addition of A4 peptide triggered an 8.7-fold or 12.1-fold increase in formation of diploids containing *Bb*.Ste2 (p=0.004) or *Hj*.Hpr2, respectively (p=0.001), together with a 19-fold increase in the negative control lacking a GPCR (p=0.013) (Fig. 4C-D).

**Fig. 4.**
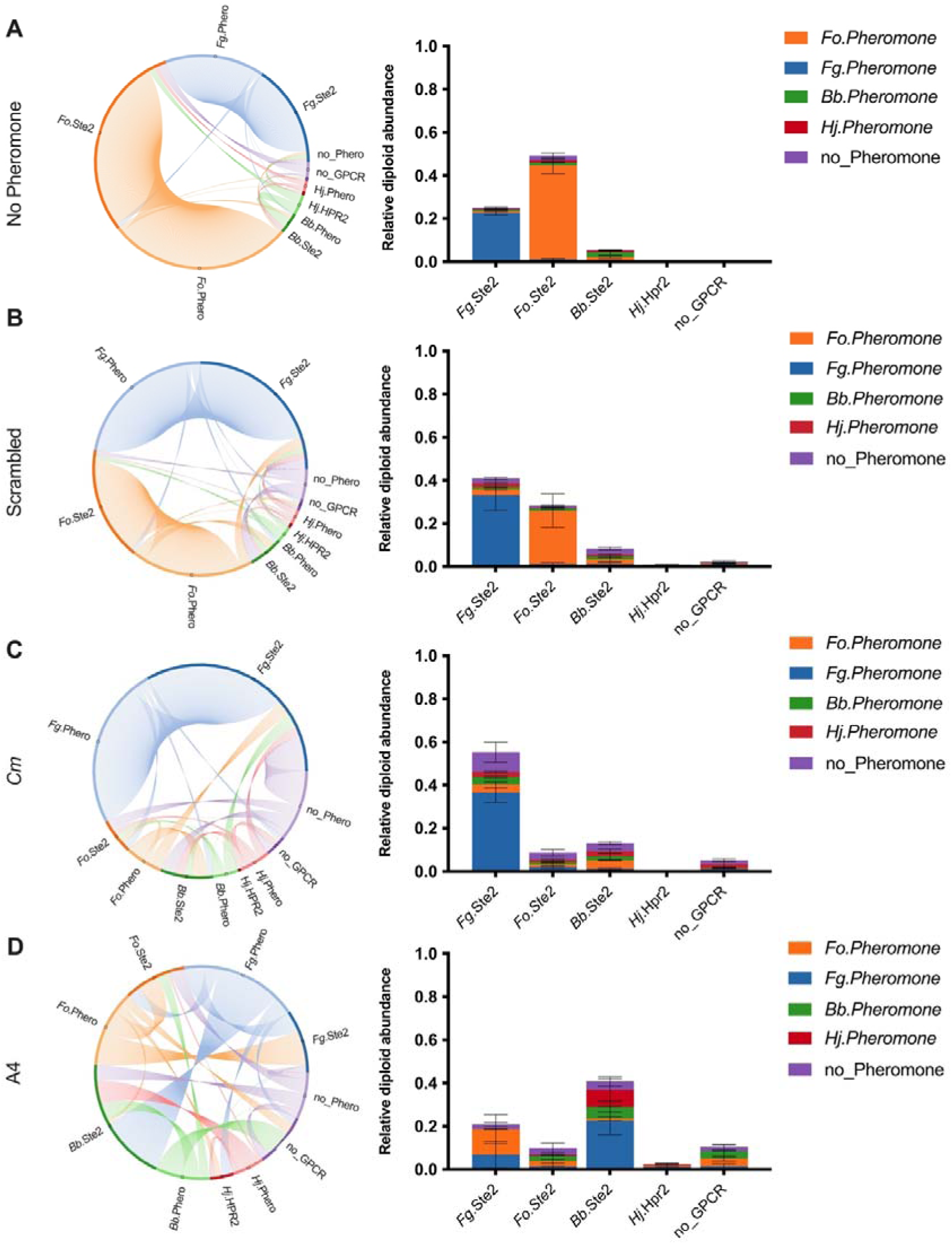
GPCR-mediated cell-cell communication in a consortium can be controlled by external pheromone supplementation. Starting from a consortium in SC media without external supplementation of pheromone in **A**, we added 10 μM of either scrambled peptide (negative control) **B**, *Cm* pheromone **C**, or A4 pheromone **D**. On the left, in the chord diagrams a dark color represents the GPCR-expressing strain while the lighter version represents the strain secreting the cognate pheromone. Orange=*F. oxysporum* (*Fo*), Blue=*F. graminearum* (*Fg*), Green=*B. bassiana* (*Bb*), Red= *H. jecorina (Hj)*, Violet=negative control with empty constructs. Every arc represents the connection between a GPCR and a pheromone strain. The size of an arc reflects the average relative abundance of a diploid cell. Relative quantification was done by comparing the Ct value of every possible combination of two barcodes with the Ct value of the conserved region flanking all barcodes. On the right, the bar plots represent the distribution of pheromone secreting strains that have successfully mated with each of the five GPCR-expressing strains. All experiments were conducted in three biological replicates.

A4 peptide supplementation reduced the presence of diploids with *Fo*.Ste2 by 0.2-fold (p=0.001), without increasing the presence of diploids containing *Fg*.Ste2. *Fg*.Ste2 and *Fg*.Pheromone interaction was reduced by 0.3-fold, however, *Fo.*Pheromone strain compensated for the final amount of diploids containing *Fg*.Ste2 (Supplementary Fig. 9A).

Furthermore, media with different nitrogen sources, which are commonly used as fertilizers, affected the relative distribution of diploid abundances. Compared to SC media with ammonium sulfate alone (SC), SC media supplemented both with ammonium sulfate and Urea (SC-AS/Urea) led to a 4-fold and 20-fold increase of diploids containing *Bb*.Ste2 and *Hj*.Hpr2, respectively (Supplementary Fig. 9B). By contrast, the presence of urea did not affect the abundance of diploids with *Fo*.Ste2 or the control strain with no_GPCR, but led to a 0.8-fold decrease in the abundance of diploids with *Fg*.Ste2 (Supplementary Fig. 9B).

In summary, YeMaP reconstructed a network of interactions between different fungal alpha pheromones and GPCRs, enabling us to test how external pheromone supplementation could interfere with GPCR-mediated cell-cell communication in fungal pathogens.

### YeMaP findings are reproduced in vivo in the phytopathogen F. oxysporum

To validate our YeMaP findings in an *in vivo* fungal system, we tested how external pheromone supplementation affects GPCR-dependent functions in the plant pathogen *F. oxysporum.* As previously reported^10^ we found that exogenous addition of 400 μM alpha pheromone from *Fo* or *Bb*, or of the synthetic agonist A4, caused a significant reduction of *F. oxysporum* microconidia germination (Supplementary Fig. 10). We next tested the chemoattractant activity of the different pheromones using an established chemotropism assay^11^. Interestingly, *Fo* and A4, the two peptides selected in the mating enrichment screen with the YeMaP platform (see Library-on-One section), induced a significant chemotropic response in *F. oxysporum* germ tubes, whereas the *Bb* pheromone and the scrambled peptide did not (Fig. 5A). Furthermore, when *F. oxysporum* microconidia were exposed to two competing chemoattractant gradients, A4 fully abolished hyphal chemotropism towards *Fo* pheromone, while *Bb* pheromone only achieved a partial reduction (Fig. 5B). Both A4 and *Fo* pheromone were able to invert directional growth of *F. oxysporum* germ tubes towards tomato root exudate (RE), while *Bb* pheromone annulled the RE response and scrambled peptide had no significant effect (Fig. 5C).

**Fig. 5.**
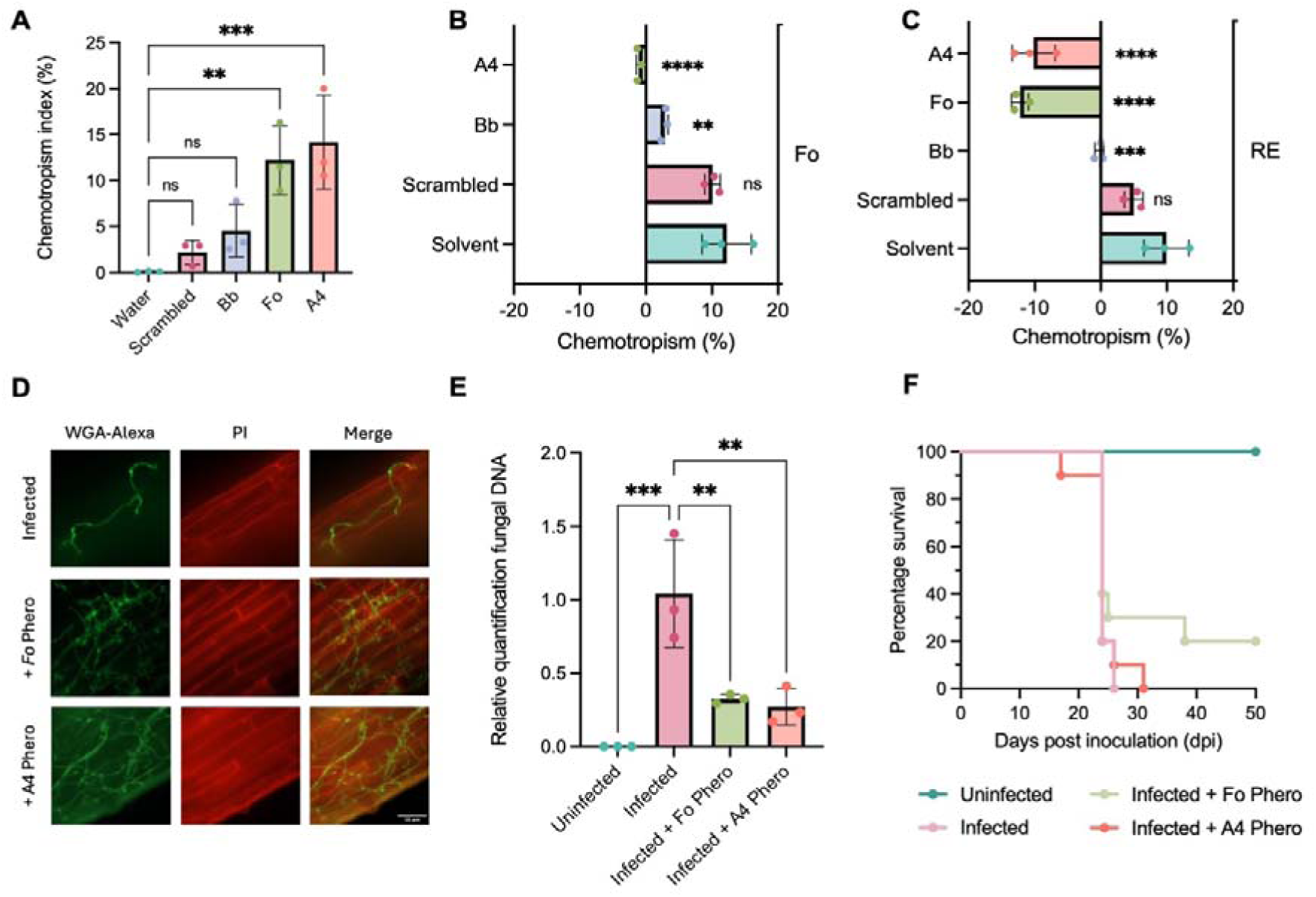
YeMaP-selected pheromones trigger *F. oxysporum* hyphal chemotropism and interfere with plant root penetration. Directed growth of *F. oxysporum* (*Fo*) germ tubes embedded in water-agar after 13□h exposure to gradients of the indicated compounds. **A** Directed growth towards gradients of 400 μM of the following pheromones: scrambled peptide (neg. control), *B. bassiana* (*Bb*), *Fo*, or A4 (Agonist 4). All pheromones were tested against the solvent (50% methanol). **B** Simultaneous exposure to competing gradients of 400 μM *Fo* pheromone, opposed to either solvent or 400 μM scrambled peptide, *Bb* pheromone or A4. **C** Simultaneous exposure to competing gradients of tomato root exudate (RE), opposed to either solvent or 400 μM scrambled peptide, *Bb* or *Fo* pheromone, or A4 peptide. **D** Fluorescence microscopy of tomato roots subjected to the indicated treatments, 3 days after inoculation with microconidia of *F. oxysporum*. Fungal cell walls were stained with WGA-Alexa Fluor 488 (green) while plant cell walls were stained with propidium iodide (red). Scale bar, 50 μm. **E** Fungal DNA in inoculated tomato roots was measured by qPCR of the *Fol4287*-specific *actine* gene using total DNA extracted 3 days after inoculation. Fungal burden was calculated using the 2^-ΔΔCt^ method and normalized to the tomato *gapdh* gene. **F** Kaplan-Meier plot showing survival of groups of 10 tomato plants (cv. Momotaro) inoculated by pipetting on top of the roots a suspension of 3.2 x 10^6^ microconidia/ml supplemented either with the solvent (infected), or with the indicated pheromone (infected + Fo phero, infected + A4 phero). Water without microconidia was used as negative control (uninfected). Data shown are from one representative experiment. The experiment was performed twice with similar results. In **A**, **B**, **C** and **E**, means represent three biological replicates. Statistical significance was determined using one-way ANOVA with Dunnett’s multiple comparison test in GraphPad Prism (✱:✱:p ≤ 0.01, ✱✱✱p□≤□0.001, ✱✱✱✱*p* ≤ 0.0001).

Given their strong chemoattractant effect, we hypothesized that *Fo* and A4 pheromone could interfere with *F. oxysporum* infection in tomato roots. While exogenous addition of *Fo* or A4 pheromone led to increased adhesion of the *F. oxysporum* germ tubes, either among themselves or to tomato roots (Supplementary Fig. 12, Fig. 5D), after careful washing of the root surface (Supplementary Fig. 11A-B) a significant reduction in fungal biomass was detected in inoculated roots treated with *Fo* or A4 pheromone compared to the untreated infected roots (Fig. 5E). Despite this short term reduction in fungal biomass within the root, no significant difference in plant survival between the treatments were observed after 50 days (Fig. 5F).

In conclusion, the *in vivo* experiments confirmed the utility of the YeMaP in the identification of bioactive pheromones and in particular an artificial agonist with improved properties on chemoattraction of *F. oxysporum* germ tubes and in reduction of fungal root penetration during early stages of infection.

## Discussion

Fungal hyphae can sense a variety of chemical cues, allowing them to grow towards nutrient sources, mating partners, or host organisms^39^. Here we established YeMaP, a synthetic yeast mating platform able to decipher fungal GPCR-pheromone interactions. We used the platform to investigate how such interactions are affected by external factors or can be manipulated through exogenous pheromone supplementation.

YeMaP exhibited an increased resolution compared to the standard designs for yeast biosensors^13,40,41^(Fig 1E-F). This improvement can be attributed to the well-documented role of the Sst2 protein in suppressing signaling noise and GPCR activation^42–44^. Accordingly, GPCR signaling did not always translate to a comparable mating efficiency. For example, from the pheromone library screen (see Library-on-One), one of the most enriched pheromones (A4) and one of the least enriched pheromones (*Cm*) for *Fo*.Ste2, resulted in being agonists for *Fo*.Ste2 (Fig. 2D and Supplementary Fig. 3C) and consequently affected *Fo*.Ste2 activity in Library-on-Library interaction networks (Fig. 4C-D).

We used YeMaP to successfully screen a library of 8,000 pheromone variants in a single test tube, and identified the *Fo* pheromone as the most enriched for the four different Ste2 GPCRs tested. Even though we observed enrichment for the cognate pheromones, the presence of a mixture of different pheromones and synthetic peptide sequences could have led to an advantage for those with higher EC_50_ values (*Fo*, A1, and A2 in *Bb*.Ste2, *Fo* in *Fg*.Ste2) or almost no effect such as A3 in *Fg*.Ste2 (Fig. 2 and Supplementary Fig. 3A). For *B. cinerea* Ste2, a GPCR that has evolved to sense a pheromone with 9 residues instead of 10, we detected the antagonistic effect of two alpha pheromones from two different fungi, *C. militaris* and *B. bassiana* (Fig. 2F-G). However, to reach an IC_50_ the antagonist pheromones needed to be added in 2,000-fold excess compared to the agonist (Fig. 2F), a situation that may not be present in natural settings. The necessity for such high concentrations of antagonist pheromone could be partially explained by the type of assay used to test the antagonist effect on a yeast biosensor. The standard yeast biosensor strain^40^ has high expression levels of the fungal GPCR and Gα subunit, besides carrying several gene knockouts (in particular *sst2*Δ) to create a highly sensitive system suitable to detect low concentrations of agonist. Such a setting may not be ideal to screen for potential pheromone antagonists. Second, the antagonistic effect of *Cm* pheromone was maintained in the mating assay (Supplementary Fig. 3D), but only at high concentrations (50 μM), suggesting that it is a weak antagonist of *Bc*.Ste2. Future studies are required to improve the selection of pheromone antagonists and validate their effect in different fungal systems.

YeMaP successfully captured the complexity of multiple GPCR-pheromone interactions and detected the effect of different environmental factors such as pH, nitrogen sources or plant peroxidase on cell-cell communication (see Library-on-Library section). This allowed us to detected the superior robustness of the *Fo*.Ste2-alpha pheromone interaction compared to those from *Fg*, *Bb*, or *Hj* (Fig. 3). This finding may be of biological relevance, since *F. oxysporum* was shown to secrete alkalinizing peptides during plant infection, thereby actively reshaping the extracellular pH and enhancing plant root colonization^45^. At the same time, plant peroxidases trigger germ tube chemotropism via *Fo*.Ste2^11^, a GPCR which also controls microconidia germination^10^ and fusaric acid production^46^. We speculate that *F. oxysporum* may have evolved a GPCR-pheromone system that remains functional across the different environments encountered during plant infection (Fig. 3, Supplementary Fig. 7A-B, Supplementary Fig. 9B). Although both *F. oxysporum* and *F. graminearum* can sense plant roots^11^ or plant exudates^47^ through their mating receptors, our results indicate that the underlying mechanisms may be functionally different. In the presence of HRP, yeast strains expressing *Fo* components were able to maintain the GPCR-pheromone communication, while those expressing *Fg* components were not (Supplementary Fig. 7A-B).

Importantly, YeMaP results with the *Fo*.GPCR correlated with real hyphal chemotropism in *F. oxysporum*. Two of the best hits selected from the YeMaP pheromone enrichment experiment against *Fo*.Ste2, *Fo* alpha pheromone and the synthetic agonist A4, exhibited a strong chemoattractant effect towards germ tubes of *F. oxysporum* (Fig. 5A-C). In contrast to YeMaP the yeast biosensor assay was not able to discriminate between pheromones capable of inducing a strong (*Fo* and A4) or a weak chemotropic effect (*Bb*) in *F. oxysporum* (Fig. 5A-C and Fig. 1E). At the same time, in the One-vs-One YeMaP experiment, we observed the promiscuity of *Bb*.Ste2 GPCR, particularly towards the *Fo* pheromone (Fig. 1C and 1E). Moreover our results indicate that *Fo* pheromone can induce germ tube chemotropism in *B. bassiana* (Supplementary Fig. 1). Previous studies have described a role of *B. bassiana* in controlling *F. oxysporum* infection, increasing tomato plant survival rate^48,49^. Further investigations are needed to determine if this mechanism could contribute to the biocontrol activity of *B. bassiana* against *F. oxysporum*.

We used YeMaP to test if external addition of pheromone interferes with signalling through a GPCR-pheromone pair of interest. Our results were in accordance with previous studies highlighting the importance of the pheromone gradient for successful mating in yeast^29,50,51^. Furthermore, YeMaP allowed us for the first time to test in a single tube the specific effect of a supplemented pheromone against four different GPCRs that all recognize pheromones with a similar backbone (Fig. 4). External pheromone supplementation in the YeMaP led to a significant reduction of diploids containing *Fo*.Ste2, indicating that it interfered with communication between a yeast cell secreting *Fo* alpha pheromone and its mating partner expressing the GPCR (Fig. 4C-D). Similarly, supplementation of Fo pheromone or A4 peptide to *F. oxysporum* microconidia enhanced hyphal adhesion (Fig. 5D), but reduced the ability of the pathogen to penetrate tomato roots (Fig. 5E). These results indicate that alpha pheromone could be secreted by *F. oxysporum* during early stage of germination^10^ and subsequently be degraded actively after plant recognition. In support of this hypothesis, analysis of previously published transcriptomics data from early infection stages of *F. oxysporum*^52^ detected upregulation of genes functioning in the pheromone response and the cell wall integrity (CWI) MAPK cascades (Supplementary Fig. 13). Additionally, the aspartyl protease Bar1, which specifically degrades *F. oxysporum* alpha pheromone^10^, was significantly upregulated (|Log2 fold change| ≥ 2.5, *p* ≤ 0.0001), while the pheromone genes did not show significant changes in transcript levels during plant infection (Supplementary Fig. 13). While external pheromone addition significantly reduced fungal entry into tomato roots, we did not observe a clear effect on plant mortality (Fig. 5F). This could be due to the fact that external pheromone was only supplemented once during initial root inoculation, and its stability and consequently action was not sufficient to provide protection during the entire duration of the infection assay (50 days).

In the future, we envision using YeMaP as a model for studying how different soil compositions and fertilizers influence cell-cell communication within fungal consortia and for developing new molecules to control fungal pathogen virulence by perturbing their natural ability to engage in cell-cell communication.

## Online Methods

### Molecular cloning for yeast engineering

#### Gene blocks and DNA parts

Gene blocks (Integrated DNA Technologies - IDT) gDOG1 to gDOG10 (Supplementary Data 1) were codon optimized for *Saccharomyces cerevisiae* with the IDT Codon Optimization tool. All oligos and gene blocks were purchased from IDT, and coding sequences were immediately preceded by a transcriptional enhancer sequence (AAAACA).

#### PCR and DNA handling

Plasmids were assembled with USER cloning^53^ to be compatible with the EasyClone- MarkerFree system for CRISPR/Cas9 engineering^54^ and adapted to the MAD-cloning strategy^55^. All plasmids, oligos, gBlocks, and parts are listed in the Supplementary Data 1.

#### Bacterial handling for plasmid construction and propagation

Plasmids were transformed (heat-shocked) in *E.coli* DH5□ strain. The competent cells were stored in glycerol at -70°C and cultivated in Terrific Broth with ampicillin (100 mg/L) at 37°C in liquid media at 300 r.p.m. or on agar plates for 16-20 hrs.

### Strains, media, and transformation

#### Strains

Strain CEN.PK2-1C and BY4741 (EUROSCARF) were used as yeast background strains. All yeast strains can be found in the Supplementary Data 1. *Beauveria bassiana* (DSM 62075) strain was obtained from the Leibniz Institute DSMZ-German Collection of Microorganisms and Cell Cultures. All the experiments with *F. oxysporum* were conducted using *Fusarium oxysporum* f. sp. *lycopersici* 4287 (*Fol4287*).

#### Media

Yeast strains were grown in Synthetic Complete (SC) media with 2% (w/v) glucose with a pH=5.6 or in a buffered media with ammonium sulfate, urea, and the citrate phosphate buffer system (0.1 x) following Prins *et al.* protocol^56^.

For *F. oxysporum* microconidia production, the wild type strain was grown for 3–4 d in liquid potato dextrose broth (PDB) at 28 LJC and 170 rpm. Germination media^10^ was used for assesing the spore germination.

Root exudate was prepared by carefully removing 2-weeks-old tomato plants from vermiculite and washing the roots to remove any adhering substrates^11^. The plants were placed in sterile water and kept at 25 °C for 30h. The collected root exudate was filtered through a 0.22-μm Millipore membrane and stored at − 20 °C until use.

#### Transformation

Except for the Yeast pheromone library (YPL1, see below), all strains were generated using the LiAc/ssDNA/PEG method^57^. All genetic cassettes were integrated into the genome of strains harboring pEDJ391 to express CRISPR-Cas9^58^. Genomic integration was achieved with the co-transformation of a *NotI* linearized fragment with homology arms and a gRNA helper vector targeting the desired EasyClone site^54^. About 1–2□μg for each plasmid or fragment of DNA was used in all chemical transformations of *S. cerevisiae.* The strain was selected in SC with 2% (w/v) glucose media lacking the appropriate amino acid or with the addition of 100 mg/L nourseothricin (Jena Bioscience, Cat.#AB-101).

#### Yeast Pheromone Library (YPL1)

YPL1 was generated by PCR amplifying two fragments F1 (of 1978 bp generated with XI-3 UP FW + DOG105) and F2 (of 961 bp generated with DOG106 + XI-3 RV) with 129 bp of homology between each other. The plasmid pDOG44 was used as a template (Supplementary Table 3). Electroporation was conducted as described by Wäneskog et al.^32^ with few modifications. The strain GEN74 was streaked on a SC-leu plate. A colony was inoculated in 60 ml of YPD and incubated at 30°C with vigorous aeration for 24 h. 40 mL of the stationary yeast culture was added to 400 ml (1/10 dilution) of fresh YPD and incubated for 3h at 30°C with vigorous aeration. 400 ml of the yeast culture were centrifuged at 4000xg for 5 minutes by first aliquoting the yeast culture in 8 falcon tubes. The YPD supernatant was discarded, and the yeast cell pellets were pooled into 1 falcon by resuspending them in a total of 50ml pre-chilled (4°C) Milli-Q H_2_O twice. The yeast cells were pelleted by centrifugation at 4000xg for 5 minutes before resuspending in 50 ml of a 1M sorbitol and 1 mM CaCl_2_ solution. The osmotically shocked yeast cells were pelleted at 4000xg for 5 minutes, and the yeast cells were resuspended in 50 ml of a 100 mM Lithium Acetate (LiAc) and 10 mM Dithiothreitol (DTT) solution. The cells in LiAc and DTT solution were incubated at 30°C with vigorous aeration for 1h, pelleted at 4000xg for 5 minutes, resuspended in 50 ml of a 1 M sorbitol and 1 mM CaCl_2_ solution, followed by another final pelleting at 4000xg for 5 minutes. The volume of the yeast cell solution was then adjusted to the total volume of 4ml, by adding the necessary volume of a 1 M sorbitol and 1 mM CaCl_2_ solution. An aliquot of the yeast solution is removed, and serial diluted by a 10-fold serial dilution. 100 μl of this diluted cell solution from dilutions 10^-5^ and 10^-6^ was plated on a YPD 2% (w/v) agar plate that was then incubated at 30°C for 48 hours before the concentration of cells (CFU/ml) in the transformation solution was enumerated via colony count. 400 μl of the yeast solution was used per transformation reaction by mixing the cell suspension with 5 µg of plasmid DNA (or any other DNA to be transformed), 5 µg of F1, 5 µg of F2, 5 µg of pESC_URA_gRNA_XI-3 plasmid and 10μl (100µg) of boiled single-stranded salmon sperm carrier DNA (ssDNA) in an Eppendorf tube. The yeast, DNA, and ssDNA mix were then transferred to an electroporation cuvette (2mm gap), and the solution was electroporated at 2.0 kV, 200 Ohm, and 25 µFD. Surviving cells and transformed cells were enumerated by serial diluting the recovered cell suspension and then plating each dilution on YPD (surviving cells) or SC -leu -ura (transformed cells). The library was expanded in SC -leu -ura for 48 hours and aliquots were stored in glycerol stock at -70°C. We avoid the introduction of a second pheromone sequence in the diploid cells, due to the potential presence of the Cas9 and gRNA plasmids, by introducing a mutation in the XI-3 site of the GPCR strains.

### Experimental procedures

#### Mating trials

##### One-on-One in a tube

MATa and MATiZ strains were inoculated from a cryo stock into a culture tube with 1 ml of SC -leu and SC -his respectively (30□°C, 250 RPM, overnight). Reaching a final OD=0.1 with a 1:1 ratio around 5 μl from a saturated MATa culture and 5 μl from a saturated MATiZ culture were combined into a culture tube with 1 ml of SC media and for 20 hours (30□°C, 250 RPM). The haploids were plated in SC -leu (MATa strains) and SC -his (MATiZ strains) in 3 plates per strain. After 20 hours the diploid cells were appropriately diluted and plated in SC -leu -his plates. The colonies on plates were enumerated after 4 days of incubation at 30 °C using the Count This app installed on an iPhone.

##### One-on-One in a deep well plate

Similarly to the One-on-One in tube assay 2 μl from a saturated MATa culture and 2 μl from a saturated MATiZ culture were combined into 250 ml of SC media and incubated at 30 °C in a shaking incubator for 20 hours (30□°C, 250 RPM). Enumeration of haploid and diploid cells was conducted in the same way.

#### Library-on-One- GPCR against the yeast pheromone library

Two full cryotubes of the yeast pheromone library (YPL1) were inoculated into a flask containing 50 ml of SC -leu. GEN87 (*Fg*.Ste2), GEN88 (*Bb*.Ste2), GEN89 (*Bc*.Ste2), and GEN90 (*Fo*.Ste2) were inoculated from the cryostock in 4 different tubes containing 4 ml of SC -his (30□°C, 250 RPM). YPL1 was co-cultured with each single receptor in a shake flask with 50 ml of SC in two biological replicates with a final OD=0.2 and a ratio of MATa 1:1 MATiZ. After 20 hours 1 ml of the co-culture was washed twice with Milli-Q water and transferred into a flask with 50 ml of SC -his -leu (30□°C, 250 RPM). The culture was left growing for 48 hours (30□°C, 250 RPM) and 5 ml of it was used to perform a genome extraction using the yeast DNA extraction kit (ThermoFisher).

#### Library-on-Library - consortia of strains with fungal pheromones and GPCRs

MATa and MAT⍺ strains were individually inoculated from a cryo stock into culture tubes with 1 ml of SC -leu and SC -his respectively (30□°C, 250 RPM, overnight). All MATa and MAT⍺ strains were mixed in two different tubes by adding the same amount of volume from the starting tubes. 50 μl of MATa cells and 50 μl of MAT⍺ cells were combined in culture tube to reach a final volume of 5 ml (SC media) and incubated at 30 °C in a shaking incubator for 20 hours (30□°C, 250 RPM). 50 μl of the SC culture were transferred to a culture tube containing SC-his-leu reaching a final volume of 5 ml. After 48 hours, genomic DNA was extracted from the SC-his-leu using the Yeast DNA extraction kit (ThermoFisher).

#### qPCR protocol - Yeast Library-on-Library

From the *Library-on-Library* experiments, 50 ng of genomic DNA was used as a template to detect the relative abundance of diploid formation. Each condition was tested in 3 biological replicates. All qPCR reactions were performed using SYBR Master Mix (ThermoFisher) on the QuantStudio 5 system.

#### Handling of synthetic pheromones

Synthetic peptides (Supplementary Table 2) were purchased by GenScript Biotech (4□mg, ≥95% purity) and dissolved in 100% DMSO to a concentration of 1000 μM, then tenfold diluted in SC media to 1000□µM (10% DMSO). A serial dilution in SC + 10% DMSO was made to achieve a concentration of 1 x 10^-11^, 1 x 10^-10^, 5 x 10^-10^, 1 x 10^-9^, 5 x 10^-9^, 1 x 10^-8^, 5 x 10^- 8^, 1 x 10^-7^, 1 x 10^-6^, 1 x 10^-5^ M for the dose-response curve.

For the *B. bassiana* and *F. oxysporum* chemotropism assay and plant infection assays the pheromones were dissolved in 50% (v/v) MeOH to reach a concentration of 4000 μM.

#### Horse Raddish peroxidase preparation (HRP)

Horse radish peroxidase (Sigma P-8375) was resuspended in phosphate-buffered saline buffer (PBS) at a concentration of 200 µM.

#### Dose-response curves and flow cytometry

Characterization of the fungal pheromones was done by conducting dose-response analyses following Jensen *et al.* protocol^20^. The same procedure was followed for studying the antagonist effect of a pheromone with one exception. The biosensor strains were first incubated with the candidate antagonist pheromone (priming), followed by the addition of the agonist pheromone after 15 minutes (competition). All data were acquired with the NovoCyte Quanteon™ (Agilent) flow cytometer. For each condition tested, four biological replicates were analyzed, with a threshold of 10,000 events per replicate.

#### Quantification of B. bassiana chemotropism

*B. bassiana* chemotropism assays were conducted as previously described with a few modifications^11^. *B. bassiana* was revived in Oat Meal Agar (OMA) for a week. Spores were collected with water/glycerol 50% and stored at 4 °C. Spores were counted using a Neubauer chamber and dissolved to reach a concentration of 10^6^ spores/ml. On the side glass microscope slides were sterilized and 3 lines were drawn on the back side of each slide: a scoring line in blue, a solvent line in blue, and a test compound in red. On the front side, 500 μl of water agar was added to create a thin layer in which 10 μl of test compound (378 μM of pheromone or water) and 10 μl of solvent (50% v/v methanol) were added in their respective lines. 15 μl of spores were added on the scoring line and spread with a cover glass slide. After 20 hours the germination was assessed on a Leica DM4000 B microscope (Leica Microsystems) equipped with a DFC300 FX camera (Leica Microsystems).

#### Quantification of F. oxysporum chemotropism

*F. oxysporum* chemotropism was conducted as previously described^11^. For each compound or combination of compounds tested, 300 hyphal tips were scored. All experiments were performed in three biological replicates.

#### Tomato plant growth conditions

Tomato seeds (*Solanum lycopersicum* cv. Moneymaker from EELM-CSIC; susceptible to *F. oxysporum f. sp. lycopersici* race 2) were surface- sterilized by immersion in 20% bleach (v/v) for 30 min and sown in moist vermiculite. The seeds and plants were maintained in a growth chamber under the following conditions: 28 LJC, 40–70% relative humidity and a photoperiod of 14 h of 36 W white light and 10 h of darkness. Plants in vermiculite were irrigated with tap water.

#### Tomato root inoculation and fluorescence microscopy of infected roots

For microscopic observation of *F*. *oxysporum* during tomato root infection, roots of 2-week-old tomato seedlings were inoculated with *F*. *oxysporum* microconidia. Freshly obtained microconidia were resuspended in sterile water at a concentration of 3.2 x 10^6^ microconidia/mL were mixed in Eppendorf tubes and mixed either with the pheromone of interest at the desired concentration or with the solvent treatment in a volume ratio 9 microconida:1 pheromone volume. For each condition tested, 3 tomato plants were placed on a water agar plate, and 5 μl of the microconidia suspension were applied on 10 different spots on each tomato root (in total 50 μl per root). After 30 minutes at room temperature the roots were dried on a paper towel, transferred to a new water agar plate and incubated 3 days at 28°C. Then the roots of 3 plants were pooled together and stained (Supplementary Fig. 11A) with 20 µg/ml propidium iodide (PI) (Sigma-Aldrich) and 10 µg/ml WGA-Alexa Fluor 488 (Invitrogen) as previously described^59^. Wide-field fluorescence imaging was performed with a Zeiss Axio Imager M2 microscope equipped with a Photometrics Evolve EMCCD camera, using the 40X oil objective.

#### Quantification of fungal DNA in tomato roots

Tomato roots inoculated with *F*. *oxysporum* microconidia and incubated 3 days as described above. Roots were washed 3 times using a water washing bottle with Driplok vapor venting (Supplementary Fig. 11B). Total DNA was extracted from tomato roots, and *F. oxysporum* biomass was measured by qPCR as previously described^60^ using primers specific for the *F. oxysporum actin* gene. For each biological replicate, roots from 3 plants were pooled for DNA extraction. Relative fungal biomass was calculated using the 2^-ΔΔCt^ method, with primers for the *Fol4287 actin* gene (FOXG_01569) normalized to the tomato *gadph* gene.

#### Tomato plant mortality assay

Tomato plant infection assays were performed as described^61^ with the following modifications. Freshly obtained microconidia were resuspended in sterile water at a concentration of 3.2 x 10^6^/mL in Eppendorf tubes and mixed either with the pheromone of interest at the desired concentration or with solvent in ratio 9:1 (v/v). For each condition tested, 3 tomato plants were placed on a water agar plate, and 10 μl of the microconidia suspension were applied onto 5 different root spots on each tomato root (in total 50 μl per root). After 30 minutes incubation at 28°C, the plant roots were dried on a paper towel and planted into minipots with moist vermiculite. Plants were transferred to a growth chamber at the conditions described above, and plant survival was recorded daily for 50 days. Plant death was diagnosed as a complete collapse of the stem, without any green parts left, accompanied by visible proliferation of the fungal mycelium on the dead tissue. The Kaplan-Meier test was used to assess statistical significance of differences in survival among groups using the log-rank test in GraphPad Prism. The infection experiment was conducted 2 times with 10 plants per condition.

### Illumina

#### Illumina Yeast Library Preparation pre and post enrichments

Following Illumina’s recommendation protocol, 50 ng of high-quality gDNA was used for a two-step PCR. PCR1 was set with 2.5 μl of gDNA, 5 μl of DOG129 (1µM), 5 μl of DOG130 (1µM), and 12.5 μl of 2x KAPA HiFi HotStart ReadyMix. A PCR clean-up was performed with AMPure XP beads (1.8 ratio) and EtOH 80%. Before performing PCR2, 1 μl of each reaction was run on a bioanalyzer DNA 1000 chip to verify the size of 395 bp. PCR2 was performed with 5 μl of DNA, 5 μl of Nextera XT Index Primer 1 (i701, i702, i703, i704, i705, i706, i707, i708, or i709), 5 μl of Nextera XT Index Primer 2 (i501), 2x KAPA HiFi HotStart ReadyMix 25 μl and 10 μl of Milli-Q water. A second PCR clean-up was performed with AMPure XP beads (1.8 ratio) and EtOH 80%. Library size validation (427 bp) was performed on a bioanalyzer DNA 1000 chip and DNA concentration was measured using Qubit 2.0. A pool of all samples was conducted to achieve a final DNA concentration of 10 nM with a proportion of 55 % YLP1 pre-enrichment and the remaining 45% with the equimolar concentration of all the remaining enriched samples.

#### Illumina Data Processing Yeast Library Preparation pre and post enrichments

The sample was sequenced with NextSeq 500 Mid Output 150 cycles with 4 technical replicates. Amplicon analysis was conducted with the standardized workflow Natrix^62^. In brief, FastQC (v0.11.9) was used to quality control the reads, and low-quality tails were removed from the reads using PRINSEQ^63^ (v0.20.4). Trimmed reads with an average Phred score of less than 25 were discarded. DADA2^64^ (v1.18.0) was used to count the number of amplicon sequence variants (ASVs) and chimera detection was done by VSEARCH^65^ (v2.15.2). DESeq2^66^ (v1.26.0) was used for variance-stabilizing transformation (VST) to normalize the read counts, and PCA and hierarchical clustering were used to detect any outliers. Since no outliers were detected, we combined the raw reads of the technical replicates and performed reads per million (RPM) normalization. Furthermore, we subtracted the RPM counts of the control starting library to allow for a better sample comparison. The data processing and ASV count analysis were done in Jupyter notebooks which can be found at https://github.com/Synthetic-Biology-Tools-for-Yeast/Yeast-Mating-Platform.

#### RNA sequencing analysis during plant infection (F. oxysporum)

Data acquisition was conducted and described in detail in a previous publication^52^. RNA-seq data have been deposited in the Gene Expression Omnibus database^67^ (Accession No. GSE243247).

### Molecular dynamics (MD) simulation

GPCR and ligand complexes were predicted with AlphaFold2 Multimer^68^ using ColabFold^69^. Receptor-ligand complexes predicted in a similar pose to *Sc*.Ste2 receptor-ligand complexes^70^ were selected. Protein preparation and minimization, system setup, MD simulations, and data analysis were performed using Maestro (Schrödinger Release 2024- 1). GPCRs were restricted to residues in the N-terminal and the transmembrane domains (1- 291). The force field of the ligands was generated with LigPrep. GPCR-ligand complexes were prepared with the Protein Preparation Wizard^71^ at pH 5.6, which included capping GPCR chain termini with neutral acetyl and methylamide groups, assignment of histidine protonation states, and whole structure minimization. *F. oxysporum* pheromone does not establish intramolecular disulfide bonds^31^, therefore the intramolecular disulfide bond was reduced for all ligands. The System Builder module was used to prepare the orthorhombic simulation box (10×10×10 Å), with the prepared GPCR-ligand complex embedded in a pre-equilibrated membrane (300K) palmitoyl-oleoyl-phosphatidylcholine (POPC) bilayer aligned with PPM3.0 of the OPM database alignment^72^, TIP3 solvent neutralized with 0.15 Na+/Cl- ions, and the OPLS4 force field^73^. Membrane model system default relaxation was used to minimize the system, and the final snapshot was used to run five independent production runs with random seeds for each GPCR-ligand complex using Desmond^74^. Simulations were run at 300 K for 300 ns and snapshots were stored at every 500-ps interval. Aggregated MD data was analyzed with Jupyter Notebook and structural images were acquired with PyMol Molecular Graphics System version 2.5.0.

Data analysis and statistical analysis

#### Flow cytometry data and gating

All flow cytometry data were extracted as FCS files and gated in FlowLogic™ v8.3 (Inivai Technologies). For the biosensors assays P*FUS1*-yEGFP-based two gates were performed. The first on alive cells, and the second to exclude a small fraction (≈ 1%) of not responding cells^20^. Fluorescence data points were derived from gated populations’ median values and median yEGFP fluorescence intensities. Normalized yEGFP intensity was obtained by normalizing it to the mean of the background.

#### Diploid frequency on plates

After the quantification of the haploid strains and diploids, the limiting haploid A was determined, and Eq. (1) was applied.

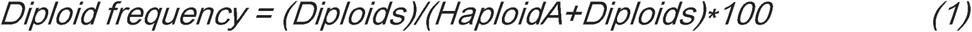

#### qPCR - diploid frequency library-on-library

The relative quantification of diploids was performed by comparing the cycle threshold (Ct) value of the conserved regions (R1 and R3) flanking the barcodes and the Ct value of the amplicon generated with two barcodes. All possible combinations of barcodes plus 3 technical replicates of the control (DOG77 + DOG78) were tested. The relative quantification was calculated using ΔCt method^75,76^.

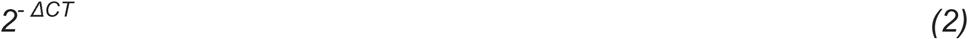

Where ΔCT = CT (barcodes) – CT (conserved region).

#### Data and statistical analysis

Data analysis, statistical analysis, and graphing were done in GraphPad Prism v10 (GraphPad Software), FlowLogic™ v8.3 (Inivai Technologies), and Jupyter Notebook. Sigmoidal dose-response curve fits were computed using nonlinear regression by the variable slope (four parameters) model in GraphPad Prism v10 (GraphPad Software), using default settings (Supplementary Data 3). The significance of the experiments was assessed by one-way or two-way ANOVA multivariate test and post-hoc analysis by Dunnett’s or Tukey’s multiple comparison test, respectively (Supplementary Data 3). All statistical tests were done in GraphPad Prism v10 (GraphPad Software) with a default 95% confidence interval (□□=□0.05) applied and multiplicity-adjusted *p* values were reported to account for all multiple comparisons within tests.

### Flow cytometry settings

#### NovoCyte Quanteon™ (Agilent) settings

Excitation was done with a blue 488□nm laser for yEGFP. yEGFP emission was detected with a 530/30□nm BP filter (471□V), while FSC and SSC were detected with a 561/14□nm BP filter (400□V). The machine ran with a core diameter of 10.1□µm (24□µl/min), two mixing cycles every well (1500□g, Acc.□= 1□s, Dur.□=□10□s), one rinse cycle every well, and a threshold for event detection at >150,000 FSC-H. NovoFlow Solution (Agilent) was used and the machine was calibrated with NovoCyte QC Particles (Agilent).

## Acknowledgements

This study is supported by grants from the Villum Foundation (00050095) to L.C. and M.K.J., the Novo Nordisk Foundation (NNF20CC0035580) (core grant), the Wenner-Gren Foundations to M.W., the Novo Nordisk Foundation (NNF23SA0084103) to T.M.F., and the Novo Nordisk Foundation Bioscience Ph.D. Program (NNF0078230) to R.T. Work in the A.D.P. lab was supported by grants PLEC2021-007777, TED2021-130262B-I00, PDC2022- 133749-I00 and PID2022-140187OB-I00 from the Spanish Ministry of Science and Innovation/State Research Agency (MICIU/AEI). M.V.A.P was supported by the PP2F_L1_01 grant co-funded by the Andalusian government and FEDER Andalucia 2021- 2027.

We thank Dr. Alvaro De Obeso-Fernández del Valle, Dr. José Mario González-Meljem, and Kinga Dulak for their technical support on the plating and the creation of the One-on-One matrix. We thank Keyan Liu for the technical support on the Illumina sequencing and Philip Tinggaard Thomsen for providing the plasmid pCfB10291. We thank Sara Lestani for the scientific discussion. Schematic figure illustrations were created with <u>BioRender.com</u>.

## Author contributions

GS, EDJ, LC, and MKJ conceived the study. GS led the investigation. GS designed and constructed all yeast strains and plasmids of this study. GS and AU performed the *B. bassiana* experiments. MM and GS performed the experiments with *F. oxysporum*. VK supported the sequencing library design. GS and MW generated the yeast pheromone library, while MJ and GS conducted the data analysis of the Illumina libraries. RT performed the Molecular Dynamics simulations. MVAP conducted transcriptomic data analysis on *F. oxysporum* during plant infection. MBS, EEHS and MD supported the investigation. GS conducted all the remaining experiments and data analysis. GS, MKJ, EDJ, and ADP wrote the manuscript. MKJ, LC, TMF, RT, MW, VK, ADP and EDJ provided the funding to enable this research.

