## Supplementary Materials for "A yeast mating platform for multiplex screening of fungal GPCR-ligand interactions"

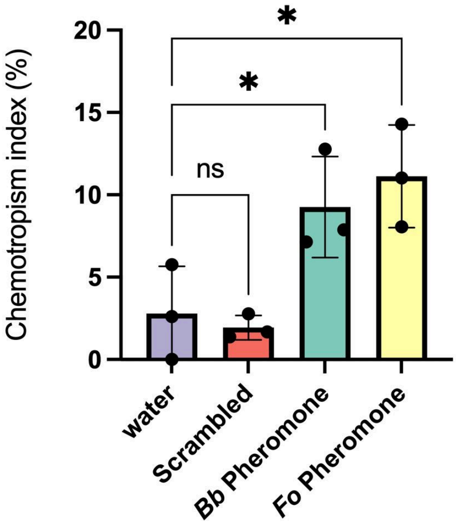


**Supplementary Figure 1.** Chemotropism of *B. bassiana* spores with different *Bb*, *Fo*, and a scrambled version of *Fo* pheromones. The assay was performed with 378 µM of pheromone and compared with the chemotropism induced by water. All points represent the average of three biological replicates; results represent the mean. Statistical significance was determined through one-way analysis of variance (ANOVA) with Dunnett's multiple comparisons (✱ $p \leq 0.05$).

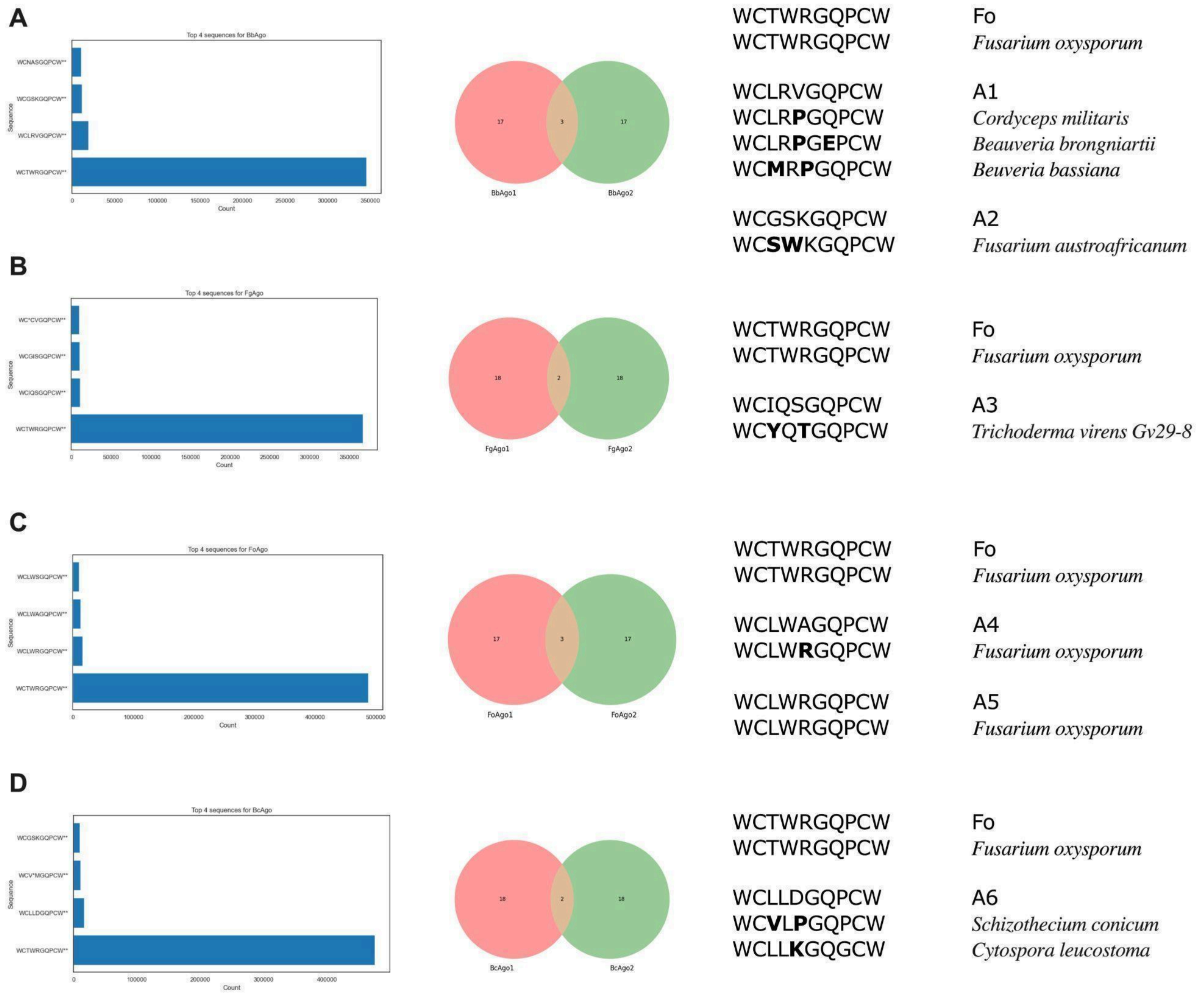


**Supplementary Figure 2.** Results of the pheromone library enrichment for *Bb*.Ste2, *Fg*.Ste2, *Fo*.Ste2 and *Bc*.Ste2. On the right are shown the counts of the top 4 pheromones enriched, and at the center, the Venn diagram with the intersection of the top 20 for the two biological replicates. The alignment of the most enriched pheromones found at the intersection of the two biological replicates is shown on the left. Except for *Fo* and A5, all the remaining pheromones exhibited some similarity to known fungal pheromones, but with one or two different residues.

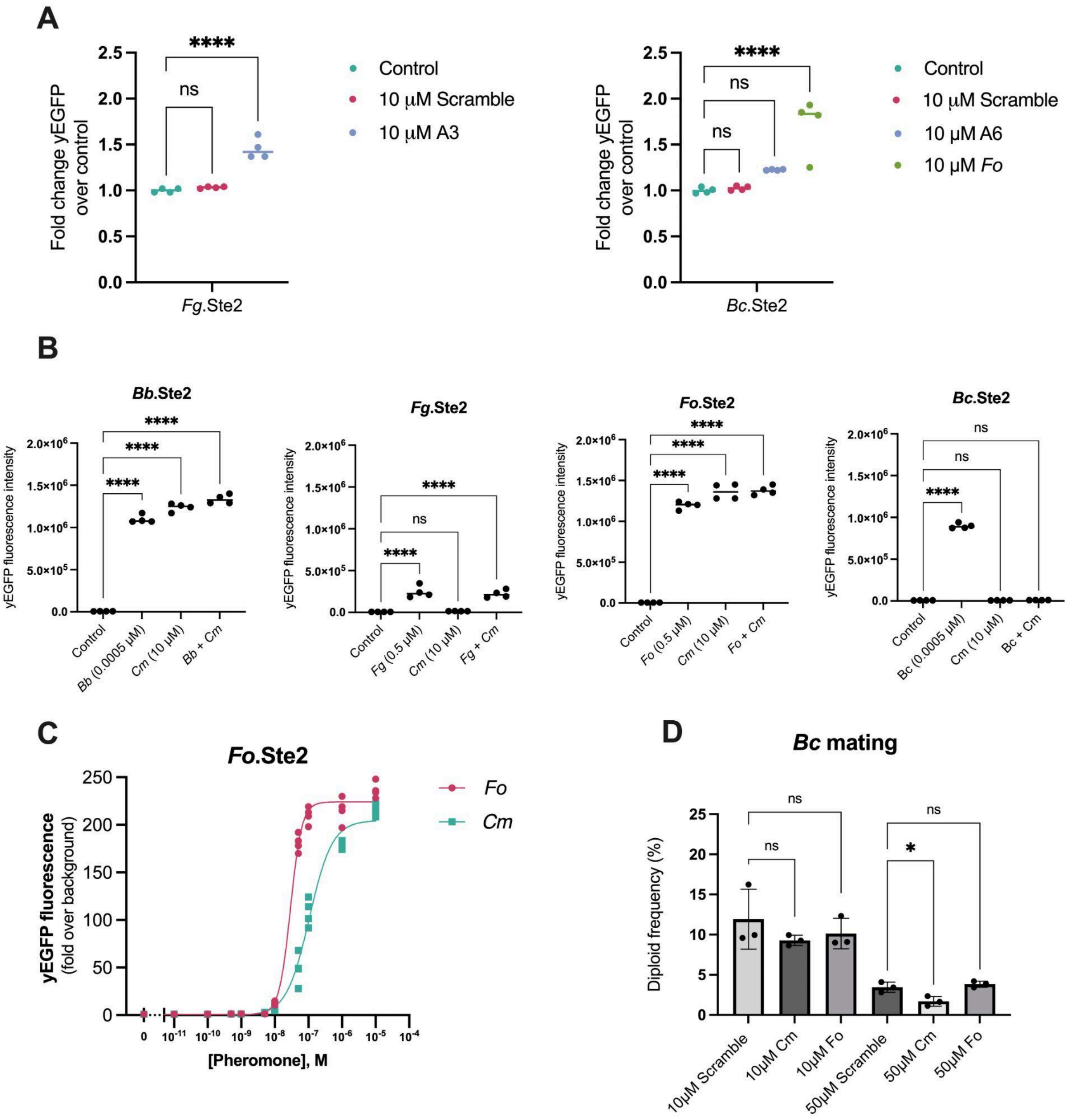


**Supplementary Figure 3. A** Fold-change activation of *Fg*.Ste2 and *Bc*.Ste2 with 10 µM of pheromone. **B** The effect of *Cm* pheromone incubated alone or in a mixture with the cognate agonist pheromone was compared to the control with no pheromone supplementation for *Bb*.Ste2, *Fo*.Ste2, *Fg*.Ste2 and *Bc*.Ste2 biosensor. **C** Dose-response curve of *Cm* pheromone on Fo.Ste2. **D** Effect of *Cm* Pheromone supplementation on diploid frequency formation with *Bc* strains. In **A**, **B**, and **C** means represent four biological replicates, and in **D** means and standard deviations represent the results of three biological replicates. In **A**, **B,** and **D** statistical significance was determined using one-way ANOVA with Dunnett's multiple comparison test in GraphPad Prism (✱$p \le 0.05$, ✱✱$p \le 0.01$, ✱✱✱ $p \le 0.001$, ✱✱✱✱$p \le 0.0001$).

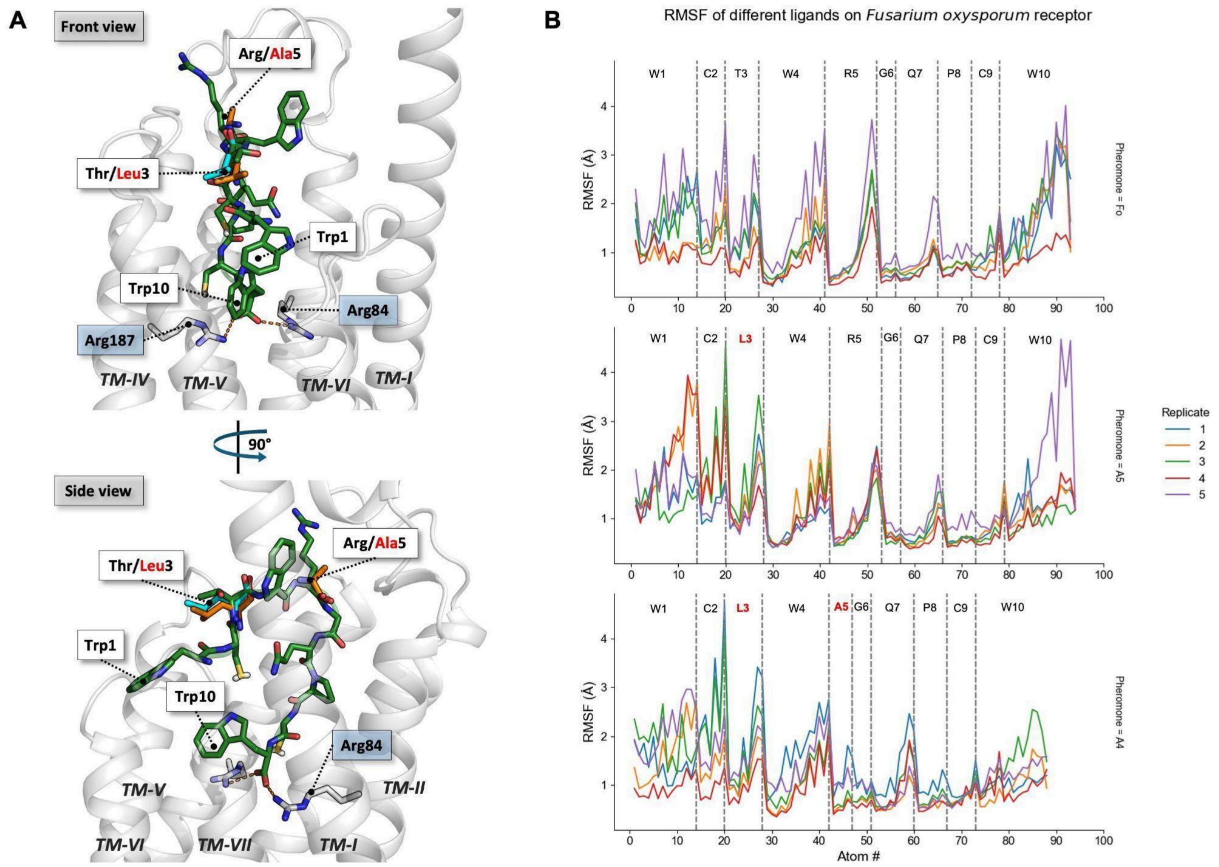


**Supplementary Figure 4. A** Proposed Binding pose of *Fo* pheromone (green) in *Fo*.Ste2. Arg84 and Arg187 interact with the carboxylic acid functional group of Trp10. A5 (cyan) and A4 (orange) pheromones deviate from the *Fo* pheromone by 1 and 2 residues, respectively. **B** Root-mean-square-fluctuation (RMSF) of non-hydrogen atoms of *Fo*, A5, and A4 pheromones inside *Fo*.Ste2 pocket. Dashed vertical lines separate the pheromone residues. Residues of A5 and A4 pheromones highlighted in red deviate from the *Fo* pheromone. Each GPCR-ligand complex prepared at pH 5.6 was submitted to 5 independent runs for 300 ns at 300 K.

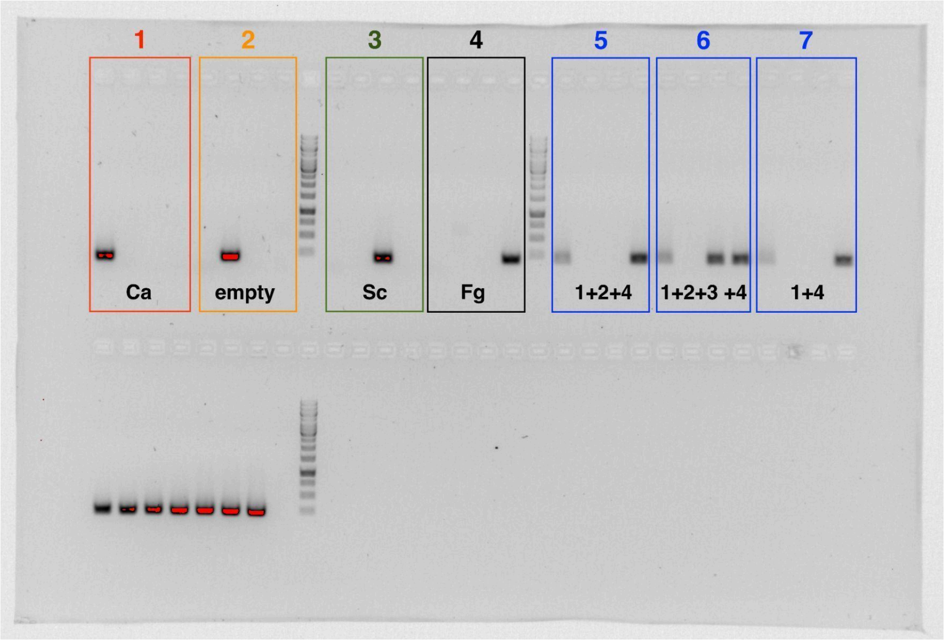


**Supplementary Figure 5.** Co-cultures (boxes 1, 2, 3, and 4) of the *Ca* GPCR and pheromone strains (GEN101 + GEN102), empty pair (GEN104 + GEN105), *Sc* pair (GEN106 + 107), and *Fg* pair (GEN108+ GEN109) with a specific amplification for their barcodes combination (in order *Ca*, empty, *Sc*, *Fg* primers). Consortia of different co-cultures of strains (boxes 5, 6, and 7). Box 5 contained the *Ca* + empty + *Fg* co-cultures, box 6 contained all four co-cultures, and box 7 contained the *Ca* + *Fg*. On the bottom, all co-cultures or consortia (1 to 7) had the R1 and R3 conserved regions flanking the two barcodes.

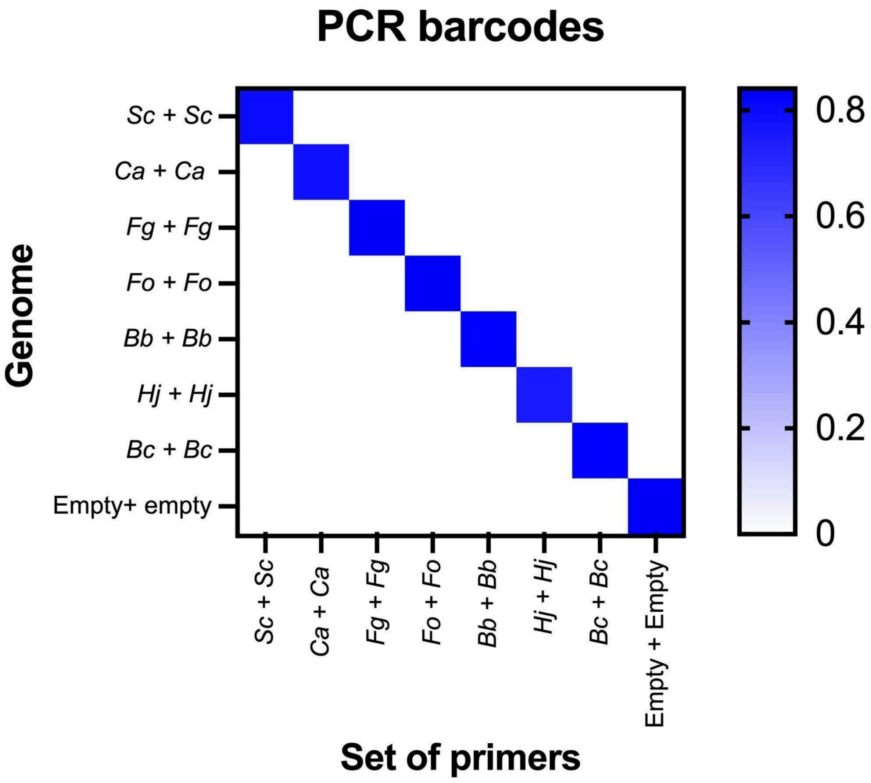


**Supplementary Figure 6.** After co-culture, the genome of the diploid cells containing two specific barcodes was used as a template for different sets of primers. We didn't observe any unspecific amplification. The proportion of diploid cells detected with different primer sets was consistent in all co-cultures (Supplementary Data 2-3).

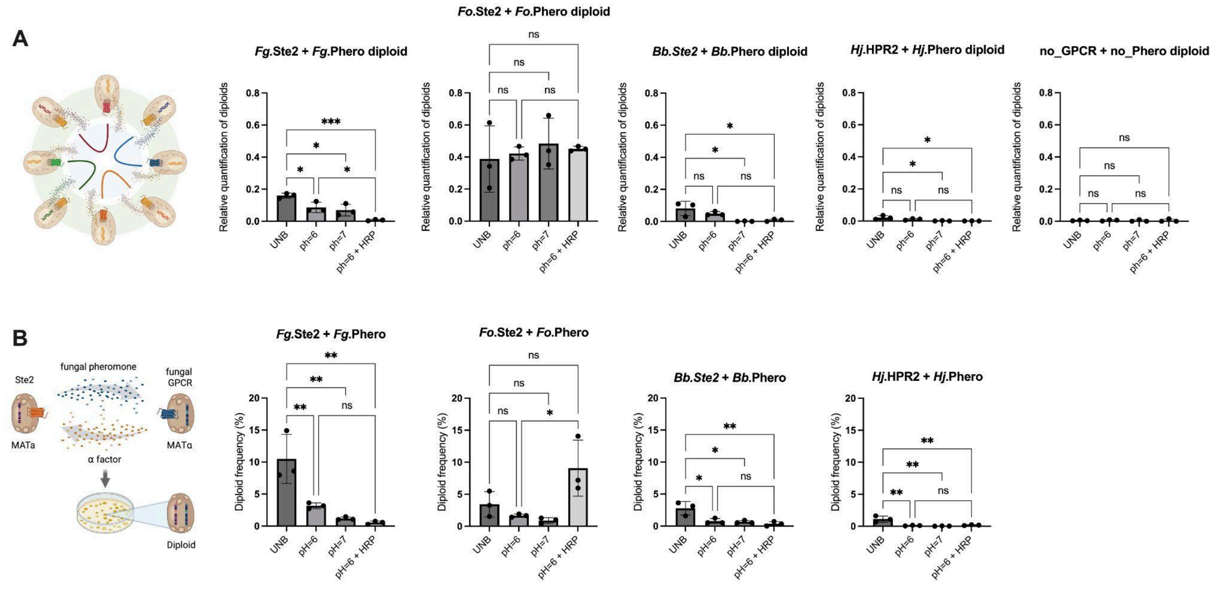


**Supplementary Figure 7.** Comparison between **A** the consortia with *Fg*, *Fo*, *Hj*, *Bb* GPCRs, and pheromone strains by adding two negative controls (empty strains) in liquid cultures (GEN108 + GEN109 + GEN114 + GEN115 + GEN110 + GEN111 + GEN116 + GEN117 + GEN104 + GEN105) and **B** the co-culture of *Fg* (GEN108 + GEN109), *Fo* (GEN114 +

GEN115), *Bb* (GEN110 + GEN111) and *Hj* (GEN116 + GEN117) strains on plates. Means and standard deviations represent the results of three biological replicates. Statistical significance was determined using one-way ANOVA with Tukey's multiple comparison test in GraphPad Prism (✱$p \leq 0.05$, ✱✱$p \leq 0.01$, ✱✱✱ $p \leq 0.001$).

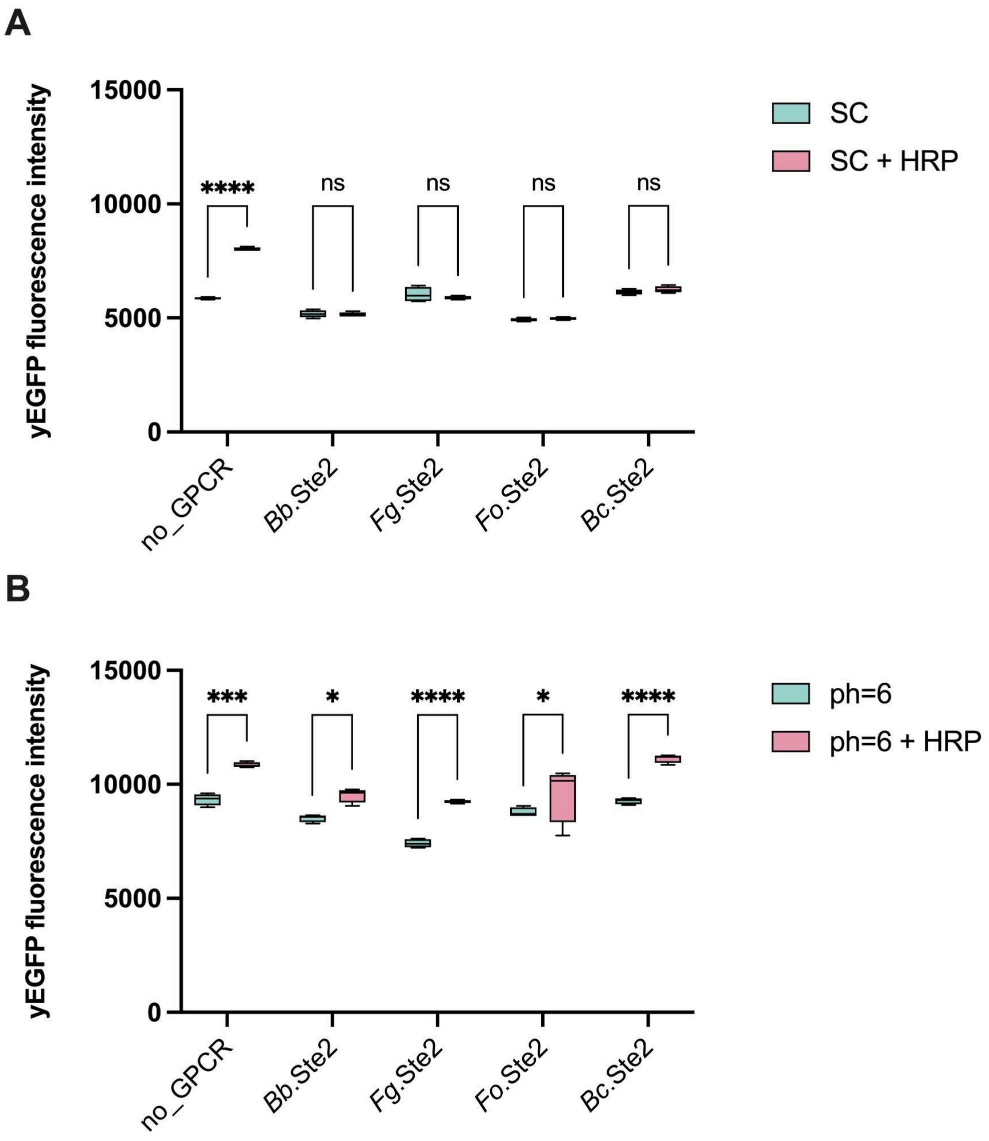


**Supplementary Figure 8.** Effect of HRP on yeast biosensors. **A** Activation of the pheromone response pathway was only observed in the background strain without a GPCR in SC media. **B** A general activation trend was observed in SC-AS/Urea pH=6 + 2 µM in all the biosensors *Bb*.Ste2 (GEN88), *Fg*.Ste2 (GEN87), *Fo*.Ste2 (GEN90) and *Bc*.Ste2 (GEN89) and the background control strain with no GPCR (CPK423). Statistical significance was determined using two-way ANOVA with Tukey's multiple comparison tests in GraphPad Prism (✱$p \leq 0.05$, ✱✱✱$p \leq 0.001$, ✱✱✱✱$p \leq 0.0001$)

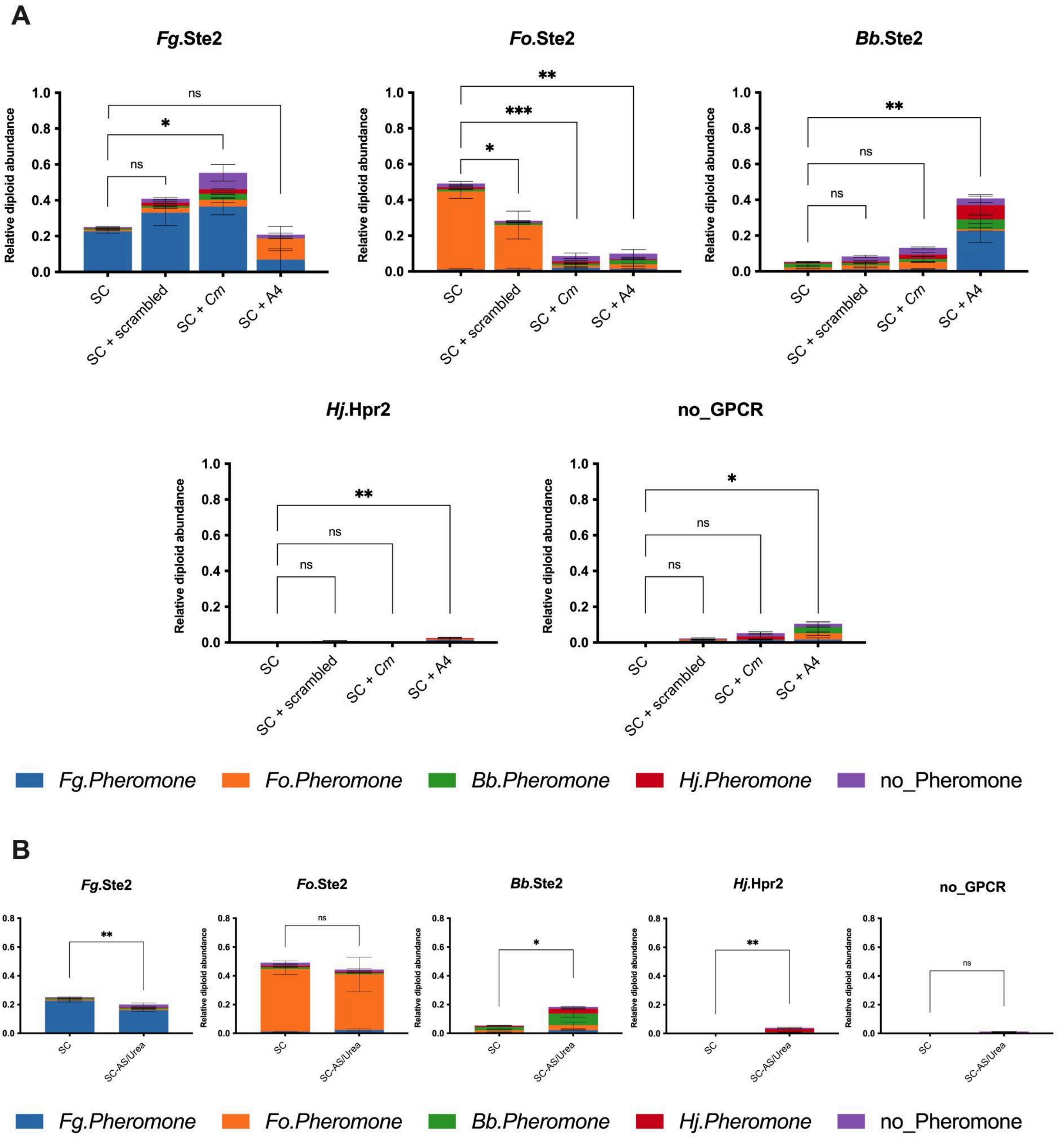


**Supplementary Figure 9. A-B** Relative diploid abundance for each fungal GPCR from consortia of *Fg*, *Fo*, *Hj*, *Bb*, negative control GPCRs, and pheromone strains (GEN108 + GEN109 + GEN114 + GEN115 + GEN110 + GEN111 + GEN116 + GEN117 + GEN104 + GEN105). **A** Diploid distribution in SC media without (SC) and with different supplementation of pheromones (SC + scrambled, SC + *Cm,* or SC + A4). All pheromones were supplemented at 10 µM. **B** Comparison of the relative diploid abundance between SC with ammonium sulfate (SC) and SC with ammonium sulfate and Urea (SC-AS/Urea). Statistical significance plotted on graphs was determined using one-way ANOVA with Dunnett's multiple comparison test in GraphPad Prism (✱$p \leq 0.05$, ✱✱$p \leq 0.01$, ✱✱✱$p \leq 0.001$) in **A,** while unpaired t-test (✱$p \leq 0.05$, ✱✱$p \leq 0.005$) was used in **B**.

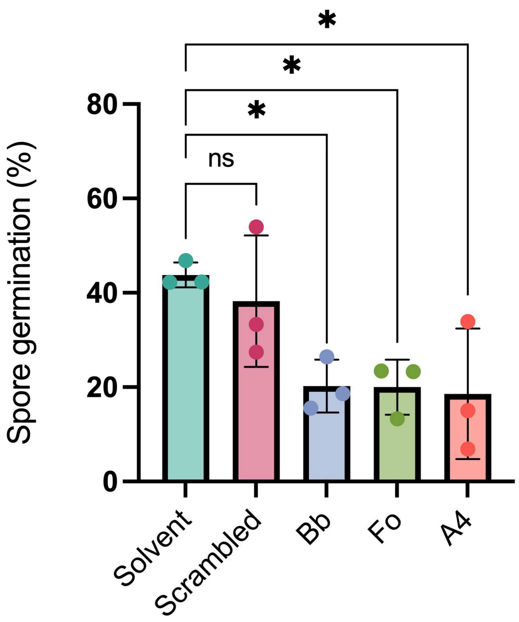


**Supplementary Figure 10.** Germination of *F. oxysporum* microconidia after 13 h of incubation in Germination Media at a cell density of 3.2 x $10^6$ microconidia/mL. The assay was performed with 400 µM of pheromone. All points represent the average of three biological replicates in which at least 300 spores were counted. Statistical significance was determined through one-way analysis of variance (ANOVA) with Dunnett's multiple comparisons (✱$p \leq 0.05$).

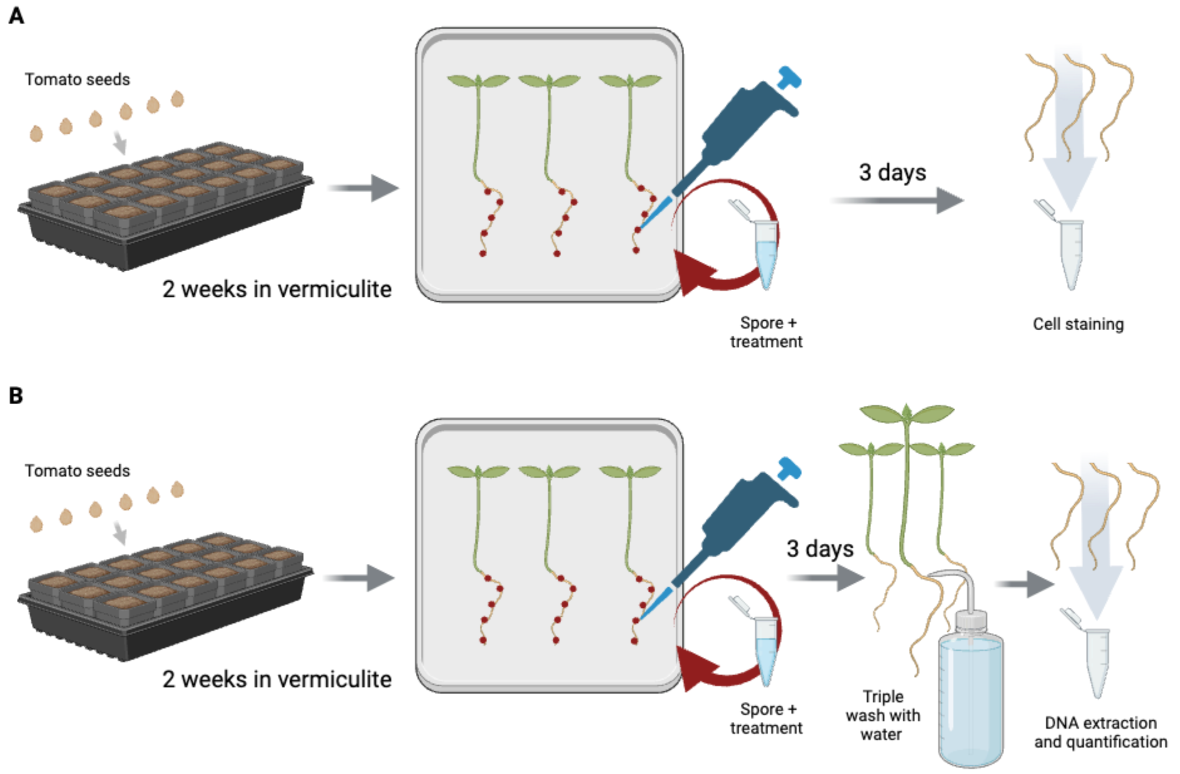


**Supplementary Figure 11.** Representation of the plant infection assay for **A** fluorescence microscopy of *F. oxysporum* infection, and **B** the relative quantification of fungal biomass in tomato roots.

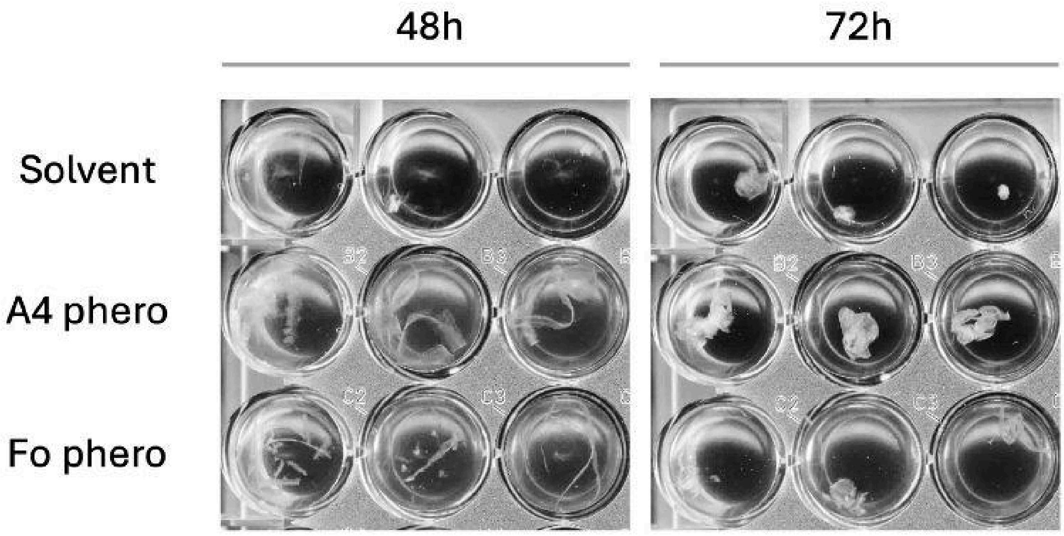


**Supplementary Figure 12.** Formation of hyphal aggregates after 48 and 72 hours in the plant root exudate media. The microtiter plate with a cell density of 2.5 x $10^6$ microconidia/mL was maintained at 28℃ and 170 rpm.

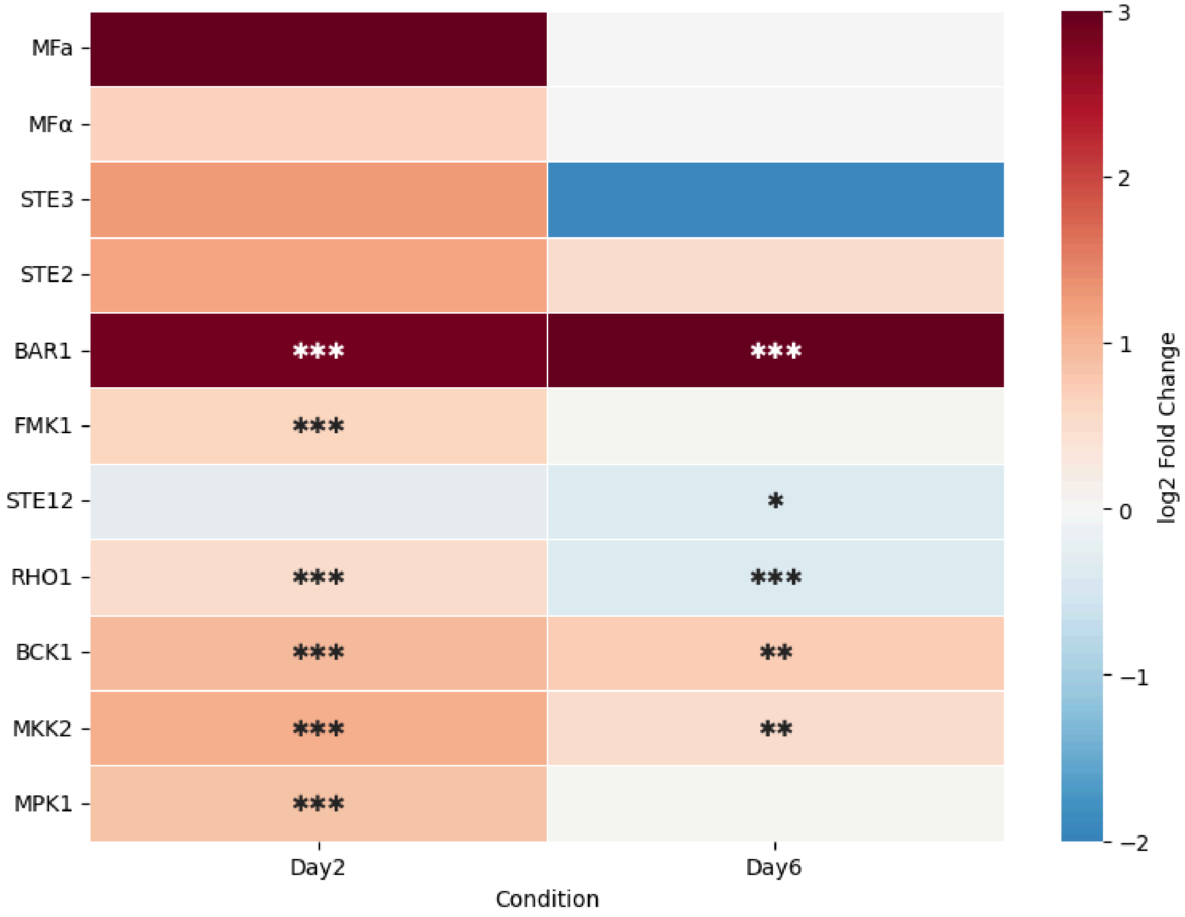


**Supplementary Figure 13.** Heatmap showing the relative expression in *F. oxysporum* after 2 and 6 days of tomato plant infection. The selected genes include pheromones, mating receptors, BAR1, and those involved in the MAPK pheromone response pathway and the cell wall integrity (CWI) MAPK cascade. The fold-change and the p-value were calculated with DESeq2 (✱$p \leq 0.05$, ✱✱$p \leq 0.01$, ✱✱✱$p \leq 0.001$).
